# Multi-omics approach combined with functional characterization of genes enlightens the formation of certain aromatic compounds in *Vitis vinifera* cv. Assyrtiko grape berries

**DOI:** 10.1101/2025.07.25.666847

**Authors:** Konstantina Leontaridou, Urska Vrhovsek, Cesare Lotti, Eirini Sarrou, Anastasia Karioti, Anthi Katsiarimpa, Konstantinos Bakasietas, Angelos K. Kanellis

**Affiliations:** Group of Biotechnology of Pharmaceutical Plants, Laboratory of Pharmacognosy, Department of Pharmaceutical Sciences, Aristotle University of Thessaloniki, 524 41 Thessaloniki, Greece; Research and Innovation Centre, Fondazione Edmund Mach, San Michele all’Adige, Italy; Institute of Plant Breeding and Genetic Resources, Hellenic Agricultural Organization - DEMETER, 57001 Thessaloniki, Greece; Laboratory of Pharmacognosy, Department of Pharmaceutical Sciences, Aristotle University of Thessaloniki, 524 41 Thessaloniki, Greece; Department of Pharmacy, University of Pisa, Via Bonanno 33, 56126 Pisa, Italy; Vine Nursery Bakasieta – VNB, 205 00 Leontio Nemeas, Korinthos, Greece

**Keywords:** Assyrtiko, multi-omics, gene functional characterization, grape berries, eugenol, phenylacetaldehyde, raspberry ketone, *Vitis vinifera*

## Abstract

Assyrtiko, a grape variety with considerable historical importance in Greece, has recently garnered global interest due to its unique citrus and floral fragrances, elevated alcohol content, and its wines created on the volcanic island of Thira (Santorini). However, grown in various regions, such as Central Greece, its aromatic characteristics may vary, prompting an investigation into the molecular mechanisms accounting for these differences. This research focused on two Assyrtiko clones— A16 and E11—grown in the Nemea, Peloponnese, which displayed distinct aromatic traits. Chemical analysis during mid-ripening indicated that clone E11 yielded significantly greater amounts of floral terpenoids, including linalool, geraniol, nerol, and α-terpineol, while A16 was rich in C compounds like (Z)-3-hexenol and (Z)-3-hexenal that are linked to green and earthy scents. Transcriptomic analysis corroborated these results: clone E11 demonstrated an upregulation of genes related to pathogen resistance and to the terpene and phenylpropanoid biosynthetic pathways, such as *DXS, MECPS, CCD1, COMT, AADC*, and *vanillin synthase*, whereas clone A16 showed increased expression of genes related to volatile thiol production and flavonoid metabolism, including *GST3* and *GST4*. Further, five key genes (RZS1, AAT, EGS1, AADC, and CYP76F14) from A16 were functionally characterized through heterologous expression in *Saccharomyces cerevisiae* and *in vitro* enzymatic assays. These findings offer new insights into the biochemical pathways determining aroma diversity in *Vitis vinifera* cv. Assyrtiko.

**Highlight:** An integrated analysis of chemical and transcriptomic profiles from two clonal variants of Vitis vinifera var. Assyrtiko elucidates the molecular basis of aroma compound biosynthesis during grape berry ripening. This study includes the cloning and functional characterization of key aroma-related enzymes: raspberry ketone synthase (*RZS1*), eugenol synthase (*EGS1*), acetyltransferase of coniferyl alcohol (AAT), 8-hydroxylinalool synthase (CYP76F14) and phenylacetaldehyde synthase (*AADC*) from Assyrtiko berries.

## INTRODUCTION

Specialized metabolism in plants generates a diverse collection of low-molecular-weight molecules, many of which derive from primary metabolism (Dixon and Dickinson, 2024). Though not firmly demanded for fundamental cellular processes, these metabolites help plants to adapt by providing resistance to biotic and abiotic stressors and mediating ecological interactions such as pollination (Ji et al., 2024). Volatile organic compounds (VOCs), a pronounced class of specialized metabolites, are predominantly crucial for plant-environment interactions, serving as defense signals against pathogens and herbivores and as attractants for pollinators (Abas et al., 2022; Pang et al., 2021). Biosynthetically, these compounds are naturally classified into terpenoids, phenolic compounds, and nitrogen-containing metabolites, among others (Dudareva et al., 2013).

In *Vitis vinifera* berries, double-sigmoidal growth patterns are followed together with complex reprogramming of primary and specialized metabolism (Kanellis and Roubelakis-Angelakis, 1993; Zenoni et al., 2010; Tornielli et al., 2023; Zenoni et al., 2023). Plant specialized metabolites, including VOCs and their non-volatile precursors, build up in coordination with ripening and are key factors of varietal aroma and wine typicity (Martin et al., 2012; Matarese et al., 2024; He et al., 2023). Although viticultural techniques, terroir (van Leeuwen et al., 2018), and fermentation processes determine the final aroma of wine, the main aromatic potential derives from chemicals biosynthesized in grapes berries during development and ripening that are often stored mostly as glycosylated precursors, which are enzymatically released during winemaking (Rapp and Mandery, 1986; Ilc et al., 2016). Major classes of grape-derived VOCs include monoterpenes (e.g. linalool, geraniol), C-norisoprenoids (e.g. β-damascenone), phenylpropanoids (e g. eugenol), lipoxygenase-derived aldehydes and alcohols (C_6_ and C_8_ compounds), furans, and volatile thiols (Boatright et al., 2011; Lin et al., 2019) with terpenoids, C_13_-norisoprenoids, and lipid derivatives being especially important in determining the aromatic profiles of many wine types (Matarese et al., 2014; Río Segade et al., 2022).

Thanks to its Mediterranean climate, varied landscape, and abundance of indigenous grapes, grapevine cultivation in Greece thrives. Among others, Assyrtiko is very famous as one of the main white grape varieties traditionally grown on the volcanic island of Thira (Santorini), which is designated as a Protected Designation of Origin (PDO) area for Assyrtiko (Robinson et al., 2012). Wines produced from Assyrtiko grapes are globally acclaimed for their distinct acidity, mineral qualities, and aromatic richness, often featuring floral and citrus characteristics (Nanou et al., 2020; Alatzas et al., 2021; Tzamourani et al., 2024;). Nevertheless, it has been observed that when Assyrtiko is grown in other areas, such as mainland Greece, or when different clonal selections are employed, its sensory profile may change, suggesting that genotype-by-environment interactions and intra-varietal genetic diversity affect its volatile composition (Nanou et al., 2025).

Albeit Assyrtiko’s viticultural and enological importance, the molecular mechanism of aroma generation in its grape berries remains poorly understood- especially the factors driving clonal variability. While enzymes implicated in the biosynthesis of major aromatic components have been functionally characterized in other plant species—such as raspberry ketone synthase in *Rubus idaeus* (Koeduka et al., 2011) and eugenol synthase in *Ocimum basilicum* (Koeduka et al., 2006; Dhar et al., 2020)—and some progress has been made in grapevine for particular pathways like wine lactone precursor biosynthesis (Ilc et al., 2017), or terpene synthesis (Martin et al., 2010; 2012), thorough study connecting genetic variation to metabolic output in particular cultivars like Assyrtiko is often lacking.

To address this knowledge gap and understand the basis of potential aromatic diversity within Assyrtiko, we conducted a comparative investigation of two clones (A16 and E11) cultivated under the same conditions in the Nemea region of the Peloponnese (Vine Nursery Bakasietas – VNB). These clones were chosen based on initial observations indicating different phenotypic and perhaps aromatic traits, with E11 exhibiting earlier ripening and better disease resistance than A16. We postulated that these clones would have substantial variations in their berry metabolomes— volatile and non-volatile—and transcriptomes, hence helping to illuminate the pathways underlying their distinctive traits. Consequently, this research utilized a comprehensive methodology that integrates comparative metabolomics (GC-MS/MS for volatile compounds, LC-MS for phenylpropanoids) alongside RNA-seq transcriptomics to thoroughly delineate the biochemical and molecular distinctions between Assyrtiko clones A16 and E11 throughout the process of berry ripening. Furthermore, investigating the biosynthetic routes of aroma-related chemicals helped us to practically verify the crucial enzymatic processes, which could be behind the observed metabolic distinctions and the general aroma profile. Particularly, we sought to functionally characterize putative grapevine enzymes involved in terpenoid (CYP76F14 - wine lactone precursor synthesis), phenylpropanoid (EGS1 - eugenol synthase; AAT - acetyltransferase), amino acid-derived volatile (AADC - phenylacetaldehyde synthase), and other related volatile pathways (RZS1 - raspberry ketone synthase).

By integrating chemical analyses, gene expression, and enzyme functional characterization, this work presents new understandings into the biochemical mechanisms underlying aroma formation in *V. vinifera* cv. Assyrtiko grape berries. Elucidating the function of enzymes central to the biosynthesis of main aroma compounds contribute to a broader understanding of the metabolic basis for aroma diversity in wine grapes and contributes towards clonal selection and the preservation of the distinctive aromatic qualities of Greek cultivars.

## Materials and methods

### Plant Material

Two clones A11 and A17 of the grapevine variety Assyrtiko, selected by Vine Nursery “VNB Bakasietas” in Nemea, Peloponnese, Greece were chosen for the present study. For each clone, 150 grape berries were randomly collected from four to five bunches at the mid-ripe stage in August 2020. Berries were sampled from three plants *per* clone, frozen immediately, and transported on dry ice to the Department of Pharmaceutical Sciences, Thessaloniki, Greece, where they were stored at −80°C for subsequent chemical analysis and RNA extraction.

### Grape Sample Preparation and Free Volatiles Extraction for GC-MS/MS Analysis

For volatile compound extraction, 2.5g of frozen grape berry powder was weighed into a pre-cooled 20mL headspace vial. The sample was mixed with 2g sodium chloride (NaCl), 0.02g ascorbic acid, 0.02g citric acid, and 2.5mL Milli-Q water to facilitate volatile release and minimize enzymatic degradation. 2-Octanol was added as an internal standard for quantitative analysis. Samples were immediately subjected to solid-phase microextraction (SPME) without further incubation.

Volatile analysis was performed using a Trace GC Ultra gas chromatograph coupled to a TSQ Quantum triple quadrupole mass spectrometer (Thermo Scientific, Italy), equipped with a CTC Combi-PAL autosampler (Zwingen, Switzerland). A 2cm Divinylbenzene/Carboxen/Polydimethylsiloxane (DVB/CAR/PDMS) SPME fiber (Supelco, 57299) was used for headspace sampling. Helium was employed as the carrier gas at a constant flow rate of 1.2mL/min. The GC inlet was operated in splitless mode with a splitless time of 4min. The transfer line and ion source were maintained at 250°C. The mass spectrometer was operated in electron ionization (EI) mode, scanning from m/z 40 to 250 with a scan time of 200ms. The GC oven temperature was programmed as follows: initial temperature of 40°C held for 4min, ramped to 250°C at 6°C/min, and held at 250°C for 5min. Data acquisition and analysis were performed using XCalibur software (Thermo Scientific, Italy). Compound identification was based on comparison of retention times with those of authentic standards (when available), and by matching mass spectra against the NIST library, supported by retention index data. Quantification was performed by normalizing peak areas to the internal standard (n-heptanol). Results were expressed as µg/L of 2-octanol equivalents. A total of 55 volatile compounds were identified and quantified, and concentrations were reported as µg of compound *per* kg of grape. Multivariate data analysis and compound visualization by grape clone were conducted using MetaboAnalyst 5.0 (Pang et al., 2021).

### Chemical Structures

Structures were drawn using RCSB PDB Chemical Sketch Tool (https://www.rcsb.org/chemical-sketch).

### RNA Isolation and Sequencing

Approximately 400 mg of grape berries were ground to a fine powder using either a mortar and pestle or a laboratory mill (IKA A11; IKA-Werke GmbH & Co. KG, Staufen, Germany) under liquid nitrogen. Total RNA was extracted using the Spectrum RNA extraction kit (Sigma-Aldrich Chemie GmbH, Schnelldorf, Germany) following the manufacturer’s protocol. RNA integrity was verified via agarose gel electrophoresis and quantified using a NanoDrop 2000 Spectrophotometer (Thermo Fisher Scientific, Wilmington, USA). Sequencing was carried out on the BGISEQ-500 platform (BGI, Shenzhen, China). Sequencing reads containing adaptors or >5% unknown bases (Ns) were filtered out. The resulting clean reads were used for transcriptome assembly.

### Transcriptome Assembly and Differential Gene Expression Analysis

Transcriptome assembly was performed using Geneious Prime® version 2020.1.2, aligned against the 12X.v2 grapevine reference genome and VCost.v3 gene annotation (Canaguier et al., 2017; https://urgi.versailles.inra.fr/Species/Vitis/Annotations). Gene expression levels were estimated in FPKM (fragments per kilobase of transcript per million mapped reads). Differentially expressed genes (DEGs) between the two clones were identified using the DESeq2 package in R, with thresholds of |log2FC| ≥ 2 and adjusted p-value < 0.05. Heatmaps of DEGs were generated using ClustVis (https://biit.cs.ut.ee/clustvis/). KEGG and GO enrichment analyses were performed using DAVID (https://david.ncifcrf.gov/summary.jsp) and KEGG tools (https://www.genome.jp/kegg/kegg1b.html), with visualization via ShinyGO v0.741 (http://bioinformatics.sdstate.edu/go/).

### Quantitative Real-Time PCR (qRT-PCR)

cDNA was synthesized from 500 ng of total RNA using a First Strand cDNA Synthesis kit. qRT-PCR was performed in 10 µL reactions using Kapa SYBR Green with cDNA diluted 1:30 on an AB 7500 Real-Time PCR system (Applied Biosystems, Carlsbad, USA). Elongation factor 1a (EF1a) was used as a reference gene, and three biological replicates were analyzed per sample. Specific primers were designed using Primer3Plus (Supplementary Table S5), and relative expression was calculated using the 2^−ΔΔCt^ method.

### Strains, Plasmids, Media, Strain Construction, and Detection for Functional Characterization of the RZS1 Gene

The *Saccharomyces cerevisiae* strain AM113 (genotype: Mata/α, GALp-(K6R)HMG2::HOX2, ura3, his3, trp1, PTDH3-HMG2(K6R)X2::leu2, PTDH3-HMG2(K6R)::HO1, ERG9/erg9, UBC7/ubc7, ssm4::SSM4, mct1/MCT1, whi2/WHI2, PTDH3-CcGGDPS1-FLO8, PTDH3-SfFDPS1-FLO1) was used for the functional characterization of the RZS1 gene. This strain is engineered for enhanced monoterpenoid biosynthesis through heterozygous deletions in ERG9, UBC7, SSM4, MCT1, and WHI2, and integration of GGDPS1 from *Cistus creticus*, FDPS1 from *Salvia fruticosa*, and two copies of a modified HMG2 variant (Božić et al., 2015). The RZS1 gene from *Vitis vinifera* was identified via a BLAST search (NCBI Accession: XM_002279390.3) as an orthologue of RZS1 from *Rubus idaeus* (GenBank ID: JN166691) (Koeduka et al., 2011). The corresponding cDNA was cloned from *V. vinifera* cv. Assyrtiko 16 using the primers: RZS1 FRW (*Bam*HI): 5’-GAGAGGATCCATGGCGGAAGTGAGCAACAA-3’ RZS1 RVS (*Xho*I): 5’-GAGACTCGAGTCATTCTCTAGCAACAACAACC-3’ PCR products were inserted into the *Bam*HI and *Xho*I restriction sites of the pUTDH plasmid. The ligation product was transformed into *E. coli* NEB 10-β competent cells, and plasmids were isolated from cultures grown in LB medium (10 g/L tryptone, 5 g/L yeast extract, 5 g/L NaCl).

Yeast transformation was performed using the lithium acetate/single-stranded carrier DNA/PEG method. Transformed *S. cerevisiae* strains harboring either pUTDH-VviRZS1 or empty pUTDH vector were grown in SD selection medium (20 g/L glucose, 1.7 g/L yeast nitrogen base, 2 g/L synthetic complete dropout mix lacking uracil). After 24 h of growth at 30°C, cultures were inoculated into 20 mL of fresh SD medium at an initial OD of 0.06. p-Hydroxybenzalacetone was added to a final concentration of 0.5 mM as the substrate, and cultures were incubated for 48 h at 30°C. Metabolites were extracted by mixing culture supernatants with an equal volume of ethyl acetate (1:1, v/v), followed by sonication and centrifugation at 2000 × g for 2 min. The supernatant was collected, and the extraction was repeated three times. Combined extracts were evaporated to dryness and resuspended in 1 mL methanol for HPLC analysis.

Analysis was performed using a Thermo Finnigan Surveyor HPLC system (Thermo Fisher Scientific, MA, USA) equipped with a Hypersil GOLD column (5 μm, 100 × 4.6 mm). A binary solvent system was used: Solvent A (water), and Solvent B (acetonitrile containing 0.05% formic acid, pH 3). The gradient elution profile was as follows: 15–60% B over 20 min, followed by 60–15% A over 11 min, at a constant flow rate of 1.0 mL/min. Raspberry ketone was detected by UV absorbance at 280 nm.

### Cloning and Functional Characterization of AAT and EGS1 Genes

Full-length sequences of AAT and EGS1 genes were obtained via BLAST searches in the *Vitis vinifera* PN40024 genome assembly, using the characterized orthologues from *Ocimum basilicum* as references. The identified *V. vinifera* gene sequences (GenBank accession numbers: XM_002272399.4 for AAT and XM_003631652.1 for EGS1) were used to design gene-specific primers with Primer3Plus (https://www.bioinformatics.nl/cgi-bin/primer3plus/primer3plus.cgi), using cDNA from *V. vinifera* cv. Assyrtiko 16 as the template.

The following primers were used for restriction enzyme-based cloning: AAT FRW (*Eco*RI): 5’-GAGAGAATTCCATGGCAGCTGATCAAGTTGT-3’ AAT RVS (*No*tI): 5’-GAGAGCGGCCGCAAACCTCCATTAGAAACTCCT-3’ EGS1 FRW (*Bam*HI): 5’-GAGAGGATCCCCATGGAGAGTGTGTTAAGTATC-3’ EGS1 RVS (*Not*I): 5’-GAGAGCGGCCGCAGCCACGGAAAGAAAGTTTTT-3’ High-fidelity PCR amplification using KAPA HiFi DNA Polymerase (Roche) enabled the successful cloning of the target genes into the pXCK-K expression vector. This vector drives N-terminal fusion of the cloned genes with DnaK and a His-tag (Kyratsous et al., 2009). The constructs were transformed into *E. coli* BL21-CodonPlus (DE3)-RIL cells (Agilent Technologies, CA, USA) for high-level protein expression. Due to poor solubility of the recombinant proteins in standard lysis buffer, 0.5 M L-arginine (pH 7.0) was added to improve solubility. Purification was performed using HisSpinTrap columns (Cytiva, Marlborough, USA), followed by cleavage of the DnaK tag with thrombin (Merck, Darmstadt, Germany). Further purification and buffer exchange were carried out using Amicon® Ultra-4 Centrifugal Filter Units (Merck, Darmstadt, Germany). Protein concentrations were determined by the Bradford assay (Bradford, 1976). For *in vitro* eugenol synthesis, reactions were set up in a final volume of 200 μl containing 100 mM MES-KOH buffer (pH 6.0), 0.5 mM NADPH, 0.5 mM coniferyl alcohol, 0.3 mM acetyl-CoA, and 5 μg of each purified protein (AAT and EGS1). Reaction mixtures were incubated at 18°C for 40 minutes and extracted with 1 mL hexane for analysis by GC–MS.

Yeast strains expressing EGS1 were similarly extracted with hexane, and the organic phase was analyzed by gas chromatography–mass spectrometry (GC–MS) using a Shimadzu QP-2020 system (Kyoto, Japan) equipped with an AOC-20i Auto Injector and GC/MS Solution software. Separation was performed on an Agilent HP-5MS fused silica column (30 m × 0.25 mm × 0.25 μm). Instrument parameters were as follows: injection temperature 260°C; interface temperature 300°C; ion source temperature 200°C; electron ionization (EI) mode at 70 eV; scan range 41–450 m/z; scan time 0.50 s. Oven temperature programs used were:

- Program A: 55–120°C at 3°C/min, 120–200°C at 4°C/min, 200– 220°C at 6°C/min, hold at 220°C for 5 min
- Program B: 60–240°C at 3°C/min

Helium was used as the carrier gas at 54.8 kPa with a split ratio of 1:30. Compound identification was based on comparison with authentic standards and spectral matching using the NIST 98 and Wiley libraries.

### Cloning and Functional Characterization of CYP76F14 in Yeast

Primers were designed to amplify the coding region of the *CYP76F14* gene from *Vitis vinifera* cv. Muscat Ottonel (GenBank accession number XM_010659734.1), excluding the N-terminal signal peptide and incorporating restriction sites for cloning. The primer sequences were as follows:

CYP76F14 FRW (*Eco*RI): 5’-AGAGGAATTCATGAGAAGAGGAAGAAAGCATG-3’

CYP76F14 RVS (*Xho*I): 5’-GAGACTCGAGTCAAACCCGTACAGGTAG-3’

The amplified product was cloned into the *S. cerevisiae* expression vector pUTDH. The resulting plasmid (pUTDH-*CYP76F14*) and the empty vector (pUTDH) were transformed into *S. cerevisiae* strain AM113. Transformants were selected on SD medium lacking uracil.

Yeast precultures harboring either pUTDH-*CYP76F14* or the empty vector were grown for 24 h at 30°C, then used to inoculate 50 mL cultures at an initial OD of 0.06. Cultures were incubated at 30°C for 72 hours, followed by metabolite extraction using hexane. Hexane extraction was performed three times with an equal volume (1:1, v/v), including 15 minutes of sonication between each extraction. Extracts were concentrated by evaporation to a final volume of 2 mL and transferred to GC–MS vials for analysis of 8-hydroxylinalool.

### Phenylacetaldehyde Synthase (AADC) Cloning, Protein Purification, in Vitro Assay, and GC–MS Analysis of Phenylacetaldehyde

The *AADC* gene was cloned based on sequence homology with the characterized *Rosa × hybrida* ‘Hoh-Jun’ gene (GenBank accession number AB305071) and compared with the *Vitis vinifera* ‘Vidal blanc’ *VvAADC* gene, as reported by Pan et al. (2012). Primers were designed for restriction enzyme cloning into the bacterial expression vector pAlex as follows:

AADC FRW (*Eco*RI): 5’-GAGAGAATTCCATGGTTGGAAGCGCTCTATT-3’

AADC RVS (*Not*I): 5’-GAGAGCGGCCGCCTTATGCAGATCACACAAACAA-3’

PCR amplification was performed using cDNA from *V. vinifera* cv. Assyrtiko 16 as template. The recombinant protein was expressed in *E. coli* BL21-CodonPlus (DE3)-RIL cells (Agilent Technologies, CA, USA), as a fusion protein containing a C-terminal GST tag and an N-terminal His tag (Panagiotidis and Silverstein, 1997). Following expression, GST was removed by digestion with Factor Xa (New England Biolabs), and the protein was purified using His SpinTrap columns (Cytiva) followed by concentration and buffer exchange using Amicon® Ultra-4 Centrifugal Filter Units (Merck, Darmstadt, Germany). Protein concentrations were determined using the Bradford assay (Bradford, 1976).

For the *in vitro* assays, 200 μL reactions were prepared containing 5 mM L-phenylalanine, 0.05 mM pyridoxal phosphate (PLP), 1 mM NADH, and 5 μg of purified protein in 0.1 M potassium phosphate buffer (pH 8.0). Control reactions were prepared identically but without added protein, using buffer in its place. Reactions were incubated at 35°C for 2 hours, followed by an additional 2-hour incubation at 30°C. Metabolites were extracted twice with 1 mL of a 1:1 ethyl acetate:hexane mixture. The combined organic extracts were analyzed by GC–MS as previously described (Sakai et al., 2007).

GC–MS analysis was performed using a Shimadzu QP-2020 system equipped with an Agilent HP-5MS fused silica capillary column (30 m × 0.25 mm × 0.25 μm). The instrument was operated under the following conditions: injection temperature, 200°C; interface temperature, 200°C; ion source temperature, 200°C; electron ionization (EI) at 70 eV; scan range, 41–450 m/z; scan time, 0.50 s. Oven temperature programming was as follows: initial hold at 50°C for 3 min; ramp to 90°C at 10°C/min; ramp to 130°C at 30°C/min; final ramp to 290°C at 40°C/min with a 3 min hold. Helium was used as the carrier gas at 53.6 kPa with a split ratio of 10:1. Phenylacetaldehyde was identified by comparison with authentic standards and mass spectra from the NIST 98 and Wiley libraries.

### Plasmid Sequencing

All cloned plasmids were validated by Sanger sequencing, performed by CeMIA SA (Larissa, Greece).

### Statistical Analysis

Statistical analyses were conducted using OriginLab (Version 2023). Pearson correlation matrices were used for network analysis in Cytoscape 3.10.0 with MetScape 3. Correlation thresholds >|0.7| were applied. KEGG IDs and HMDB accessions were used for gene and metabolite mapping, respectively.

## Results

### Chemical analysis of grapes from Assyrtiko clones A16 and E11

The Assyrtiko clone A16 was chosen for its intense aroma, which is similar to that of Assyrtiko grapes cultivated in Santorini, where the volcanic soil contributes a distinctly characteristic bouquet. In contrast, clone E11 is an early-ripening variant that displays phenotypical characteristics, including smaller berry size and thicker skin, resulting in enhanced resistance to pathogens. To define the distinct aromatic and chemical profiles of A16 and E11, grape berries were collected and subjected to GC-MS/MS and LC-MS analyses. A total of 55 volatile compounds (expressed in μg/kg fresh weight) were identified by GC-MS/MS (Supplementary Table S1). Principal component analysis (PCA) based on volatile profiles showed clear separation of the two clones: three biological replicates of E11 grouped closely together, reflecting high reproducibility, and distinct separation from A16 replicates (PC1: 97.7%, PC3: 0.7%; Fig. 1A). Hierarchical clustering further grouped the biological replicates of each clone into separate clades (Fig. 1B).

**Figure 1.**
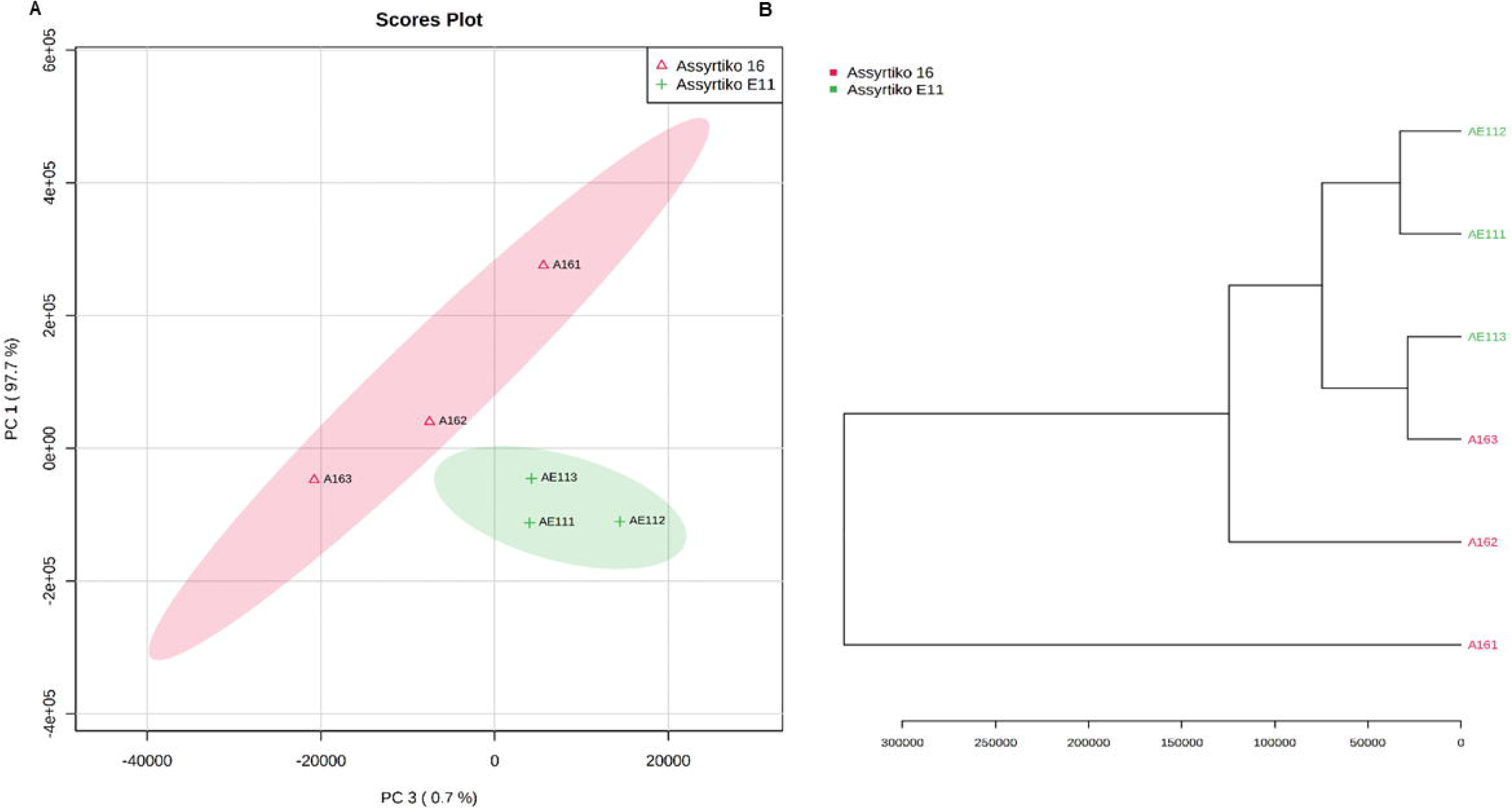
Principal component analysis (PCA) and hierarchical clustering of volatile compounds identified by GC-MS analysis of grape berries from Assyrtiko clones A16 and E11. (A) PCA plot showing separation of three biological replicates per clone based on chemical profiles. (B) Hierarchical clustering dendrogram confirming replicate grouping by clone.

Pearson’s correlation heatmap was generated (Fig. 2A) to examine the linkages between metabolites. While nerol was strongly associated with geraniol, linalool, and cis-3-hexanol, linalool had high positive relationships with nonanol, trans-2-hexenol, and trans-2-hexenoic acid. β-Ionone was co-expressed with (E,Z)-2,6-nonadienal, geranyl acetone, and trans-2-nonenal; 1-octen-3-one was related with β-damascenone, furfural, and 2,4-heptadienal. Hierarchical clustering of metabolites showed that the A16 clone had higher levels of β-damascenone (a C_13_ norisoprenoid), C_7_ and C_8_ alcohols and aldehydes; E11 had higher concentrations of terpenes, β-ionone, and C_5_ and C_6_ alcohols and aldehydes (Fig. 2B). Particularly rich in 1-octen-3-ol, terpinolene, linalool oxide C, benzyl alcohol, and furfural, A16 showed higher concentrations of geraniol, linalool, nerol, acetic acid, trans-2-hexenol, 2,6-nonadienal, 1-pentanol, and xylene, whereas E11 showed greater amounts. A two-tailed t-test (p < 0. 05) revealed two compounds that differed substantially between the clones (Supplementary Fig. S1): 1-octen-3-one, a ketone associated with earthy, mushroom-like, and metallic scents (Mosciano, 2001) was substantially more concentrated in A16; linalool, a monoterpene characterized by citrus, flowery, and rose-like odors (Mosciano, 1996), was markedly more abundant in E11.

**Figure 2.**
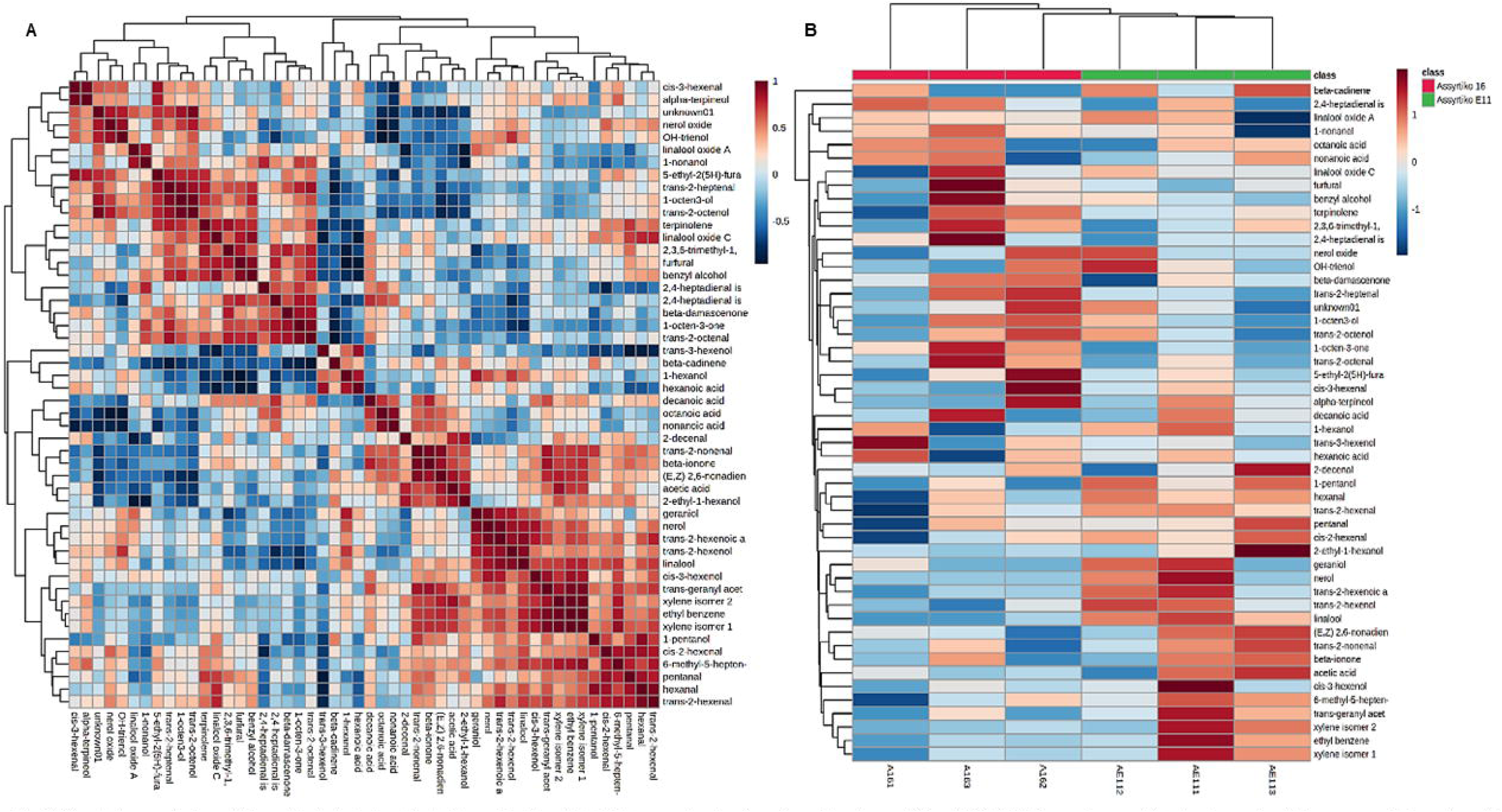
Correlation analysis and hierarchical clustering of volatile metabolites detected in grape berries from Assyrtiko clones A16 and E11. (A) Pearson’s correlation heatmap of volatile compounds based on GC-MS data (distance measure: Pearson r). (B) Hierarchical clustering heatmap of normalized metabolite levels (distance: Euclidean, clustering method: single linkage, grouped by average means).

To uncover the phenylpropanoid metabolites present in the two clones of Assyrtiko grape berries, a targeted LC-MS analysis detected 34 phenolic compounds across the two clones (Supplementary Table S2). PCA of these data separated A16 and E11 replicates (Component 1: 34.5%, Component 2: 21.6%; Fig. 3A), and hierarchical clustering confirmed distinct groupings (Fig. 3B). Although no compounds reached statistical significance between clones by univariate analysis, distinct patterns emerged based on metabolite accumulation, as visualized by a Pearson’s correlation heatmap (Supplementary Fig. S2A). Two metabolite clusters were evident: the first group included catechin, epicatechin, procyanidin, gallic acid, rutin, and p-hydroxybenzoic acid and the second contained kaempferol, ellagic acid, quercetin, and phlorizin, as highlighted by their strong correlation with p-hydroxybenzoic acid (Supplementary Fig. S2B). In both clones, epicatechin and t-coutaric acid were the most frequent polyphenols, followed by gallocatechin and catechin. While clone E11 had greater amounts of epicatechin, catechin, procyanidin, kaempferol, luteolin, and ellagic acid, clone A16 collected more t-coutaric acid, gallocatechin, and epigallocatechin. (Supplementary Fig. S2C) shows how Random Forest analysis confirmed the identification of distinguishable phenolic compounds across clones.

**Figure 3.**
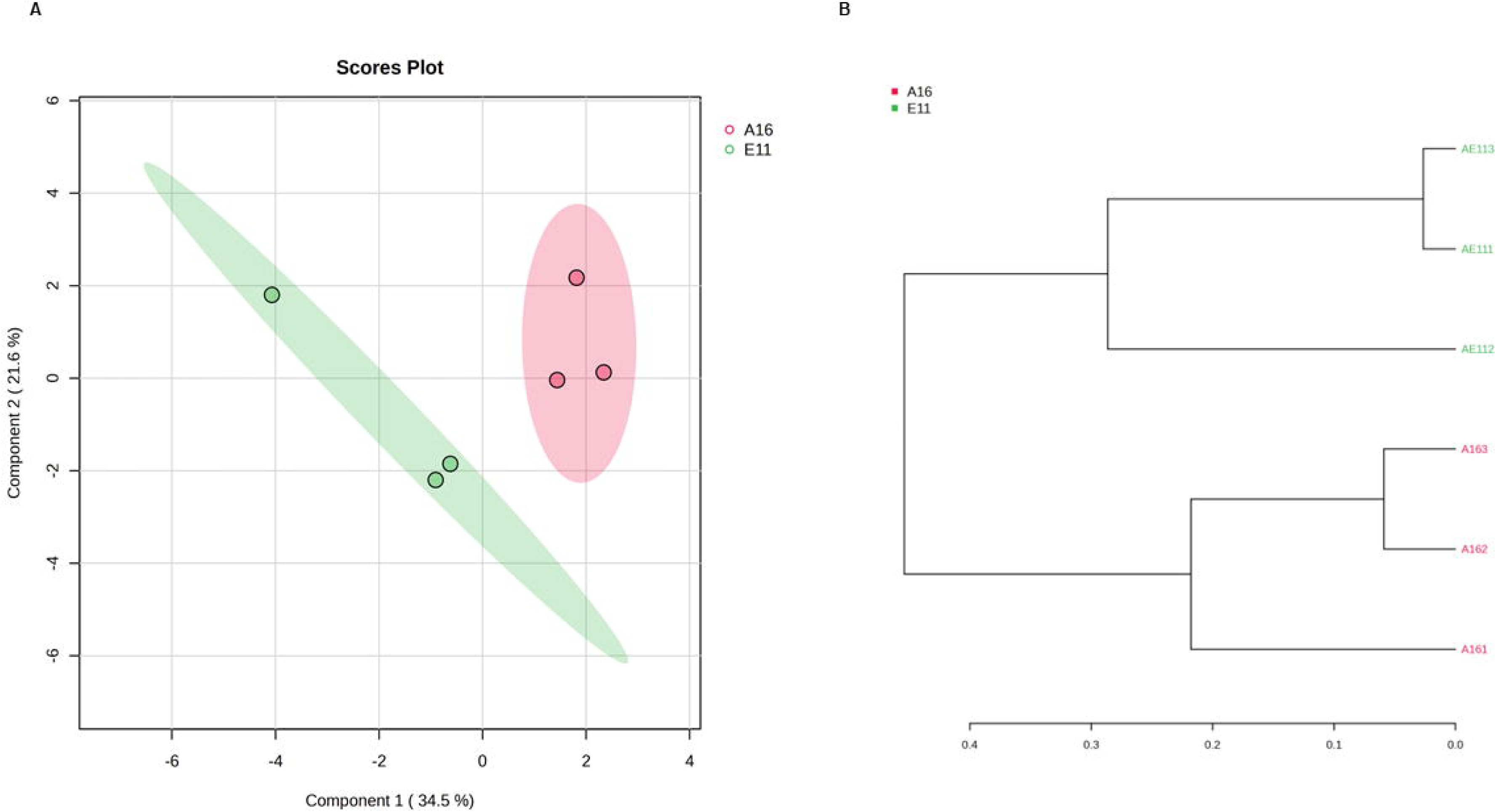
Principal component analysis (PCA) and hierarchical clustering of phenolic compounds identified by LC-MS analysis of grape berries from Assyrtiko clones A16 and E11. (A) PCA plot showing separation of biological replicates based on phenolic profiles. (B) Hierarchical clustering dendrogram illustrating clear grouping of replicates according to clone.

### Transcriptomic Analysis

Total RNA was extracted in triplicate from two clones of Assyrtiko (A16 and E11) using grape berries identical to those subjected to chemical analysis. After adapter trimming and quality filtering, RNA sequencing yielded approximately 80–97 million clean reads per sample (100 bp in length). The average GC content was ∼45%, and over 95% of bases had a Q20 score, confirming high data quality. Reads were mapped to the *Vitis vinifera* reference genome (12xV2) (Canaguier et al., 2017) using Geneious Prime, which was also used to quantify transcript abundance. A total of 42,413 transcripts were assembled, with alignment rates ranging from 94.08% to 96.71% (Supplementary Table S3).

### Differential Gene Expression Analysis and Functional Annotation of DEGs

Differential expression analysis was conducted using DESeq2. Genes with an adjusted p-value < 0.05 and |log fold change| > 2 were considered differentially expressed (DEGs). PCA of transcriptomic profiles revealed clear separation between clones, with PC1 and PC2 accounting for 33.1% and 19.86% of the variance, respectively (Fig. 4A). Replicates clustered tightly within each clone, supporting data consistency. The top 80 DEGs with the highest |log fold change| clearly distinguished the two clones (Fig. 4B). Zinc finger proteins, importins, and receptor-like kinases were upregulated in clone A16, whereas ankyrin repeats, various transporters, dehydrogenases, and RNA polymerases were more abundant in E11.

**Figure 4.**
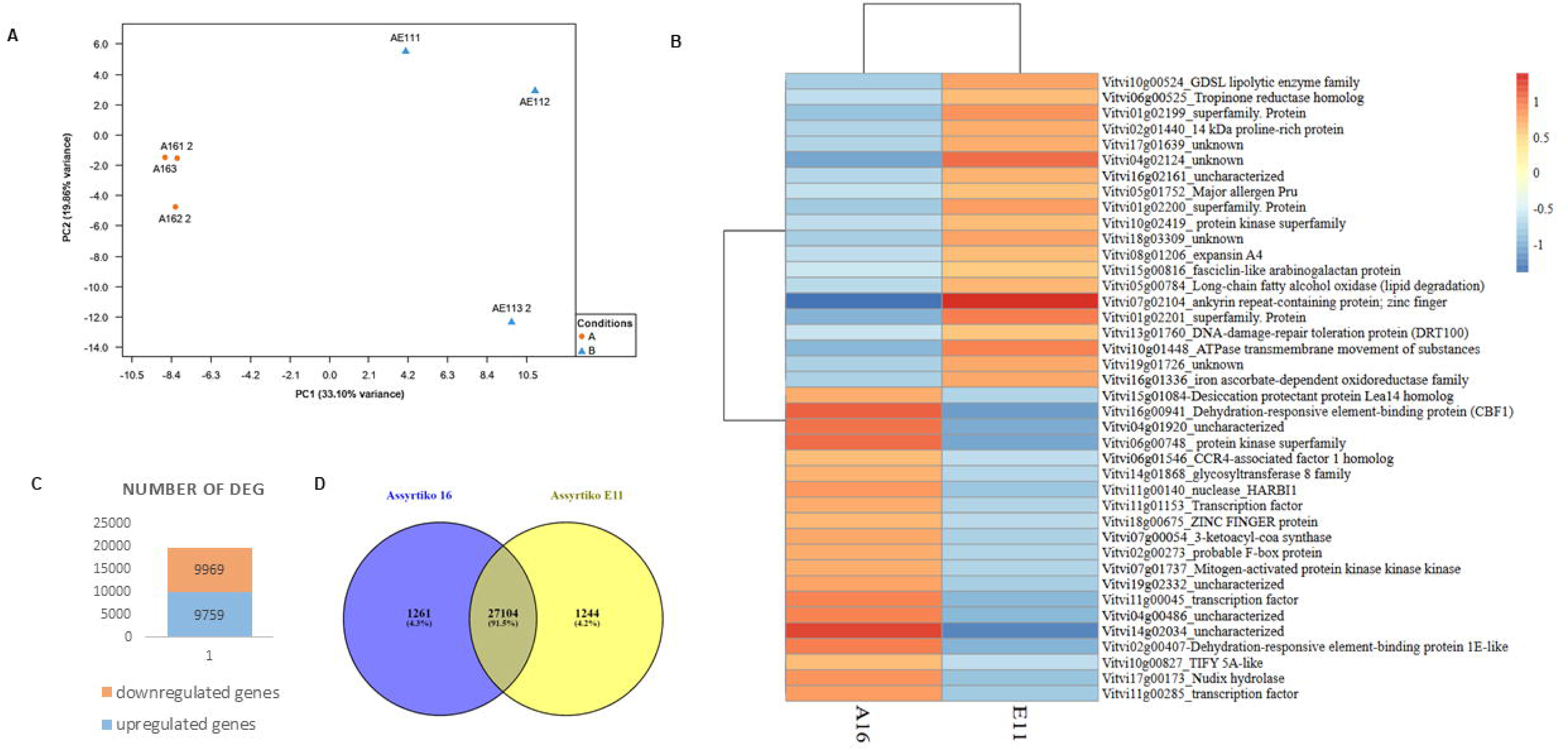
A. Principal component analysis (PCA) of differentially expressed genes (DEGs) between Assyrtiko clones A16 and E11. B. Heatmap of the 40 most highly differentiated genes (highest |log fold change| values) between the two clones, based on mean fragments per kilobase of transcript per million mapped reads (FPKM) from three biological replicates. Original values are ln(x + 1)-transformed. Rows are centered; Pareto scaling is applied to rows. Both rows and columns are clustered using correlation distance and average linkage. The heatmap was generated using the ClustVis online tool. C. Total number of the up and down-regulated genes of the Differentially Expressed Genes (DEGs). between two clones of Assyrtiko, A16 and E11. D. Venn diagram of the expressed transcripts with |log2fold change|>2, highlighting the commonly and uniquely expressed genes in A16 and E11 Assyrtiko clones with strict filter.

In total, 19,728 DEGs were identified, comprising 9,759 upregulated and 9,969 downregulated genes in A16 vs. E11 (Fig. 4C), but when applying a stricter filter (adjusted p-value < 0.01 or |log fold change| > 2) only 4.3% for A16 clone and 4.2% of the genes in clone AE11 were uniquely expressed (Fig. 4D). Genes with the highest differentiation between the two clones, that showed overexpression in E11 were an ankyrin repeat-containing protein (Vitvi07g02104), expansin 4 (Vitvo08g01206) and long-chain fatty alcohol oxidase for ω-oxidase, in lipid degradation (Vitvi05g00784), whereas in A16 clone a nuclease (Vitvi11g00140), two dehydration responsive element - binding proteins (Vitvi16g00941, Vitvi02g00407), glycosyltransferase (Vitvi14g01868) and Mitogen-activated protein kinase (Vitvi07g01737) had higher expression. Among the DEGs, 264 transcription factors (TFs) were annotated using PlantTFDB, including 147 upregulated and 117 downregulated TFs, spanning 27 families (Supplementary Fig 3). Notably enriched families in the upregulated group of transcription factors included NAC, MYB, ERF, bZIP, and bHLH. Certain families, such as C2H2 (linked to development), ZF-HD (stress-responsive), and HD-ZIP (regulating organ development and hormone response), were exclusive to the upregulated genes. Conversely, the G2-like TF family, part of the GARP superfamily involved in gene regulation and chloroplast development, was unique to the downregulated group (Supplementary Fig 3B).

GO enrichment analysis of the DEGs was performed using ShinyGO to investigate functional differences between clones (Supplementary Fig. S4). The three categories for genes were cellular component, molecular function, and biological process. Numerous genes that were upregulated were found in the cytosol and endomembrane system. The molecular functions of phosphatase, glycosyltransferase, and transporter activity were enhanced. With particular enrichment in sphingoid metabolism, coenzyme metabolism, endosomal transport, and ammonium ion processing, important biological processes included small molecule metabolism, the biosynthesis of organonitrogen compounds, and carbohydrate metabolism.

RNA/chromatin binding and catalysis were the functions of downregulated genes, which were enriched in plastid and nucleus-related components. Their functions in biology included nuclear import, transcriptional regulation, RNA processing and modification, and the organization of organelles and chromosomes.

KEGG pathway analysis revealed 136 enriched pathways among DEGs (Supplementary Fig. S5). The majority of the upregulated genes were linked to ribosomal activity, carbon metabolism, amino acid biosynthesis, and the biosynthesis of specialized metabolites. Additional enriched pathways included aminoacyl-tRNA biosynthesis, cysteine and methionine metabolism, endocytosis, and plant hormone signal transduction. Spliceosome assembly, mRNA surveillance, RNA degradation, and nucleocytoplasmic transport were all linked to downregulated genes. Both up- and down-regulated groups shared genes related to the biosynthesis of aromatic compounds. Downregulated genes were more likely to have flavonoid biosynthesis, monoterpenoid metabolism, linoleic acid metabolism, RNA replication, and spliceosome-related pathways, whereas upregulated genes were more likely to have phenylpropanoid, sesquiterpenoid, diterpenoid, α-linolenic acid metabolism, and carotenoid biosynthesis.

### Transcripts Related to Aroma Biosynthesis

KEGG pathway analysis (Supplementary Fig. S5) indicated significant differences in aroma-related biosynthetic pathways between clones A16 and E11. Based on this, key genes involved in aroma formation in grape berries were selected from the DEGs identified in the A16 vs. E11 comparison, and their normalized FPKM values were visualized in a heatmap (Fig. 5). Several carotenoid cleavage dioxygenases (CCDs) were upregulated in clone E11. Likewise, several genes involved in the phenylpropanoid and MEP (methylerythritol phosphate) pathways, along with a linalool–nerolidol synthase gene, TPS54, exhibited higher expression in E11, while another linalool–nerolidol synthase gene, TPS56, was overexpressed in clone A16 (Fig. 5). In contrast, the LOXA and LOXO genes, associated with the lipoxygenase pathway, were overexpressed in A16, as well as eugenol synthase 1 (EGS1), raspberry ketone synthase (RZS1), aromatic amino acid decarboxylase (AADC), related with phenylacetaldehyde synthesis and glutathione-S-transferases, implicated with thiol production. Many transcription factors, that regulate the aroma biosynthesis, such as MYC jasmonic acid 3 (VIT_02s0012g01320), Ethylene responsive element binding factor (VIT_16s0013g00890), WRKY DNA-binding protein 70 (VIT_13s0067g03140), Zinc finger (C2H2 type) protein (ZAT11) (VIT_06s0004g04180) and Salt tolerance zinc finger (VIT_18s0001g09230) showed higher expression in A16 clone.

**Figure 5.**
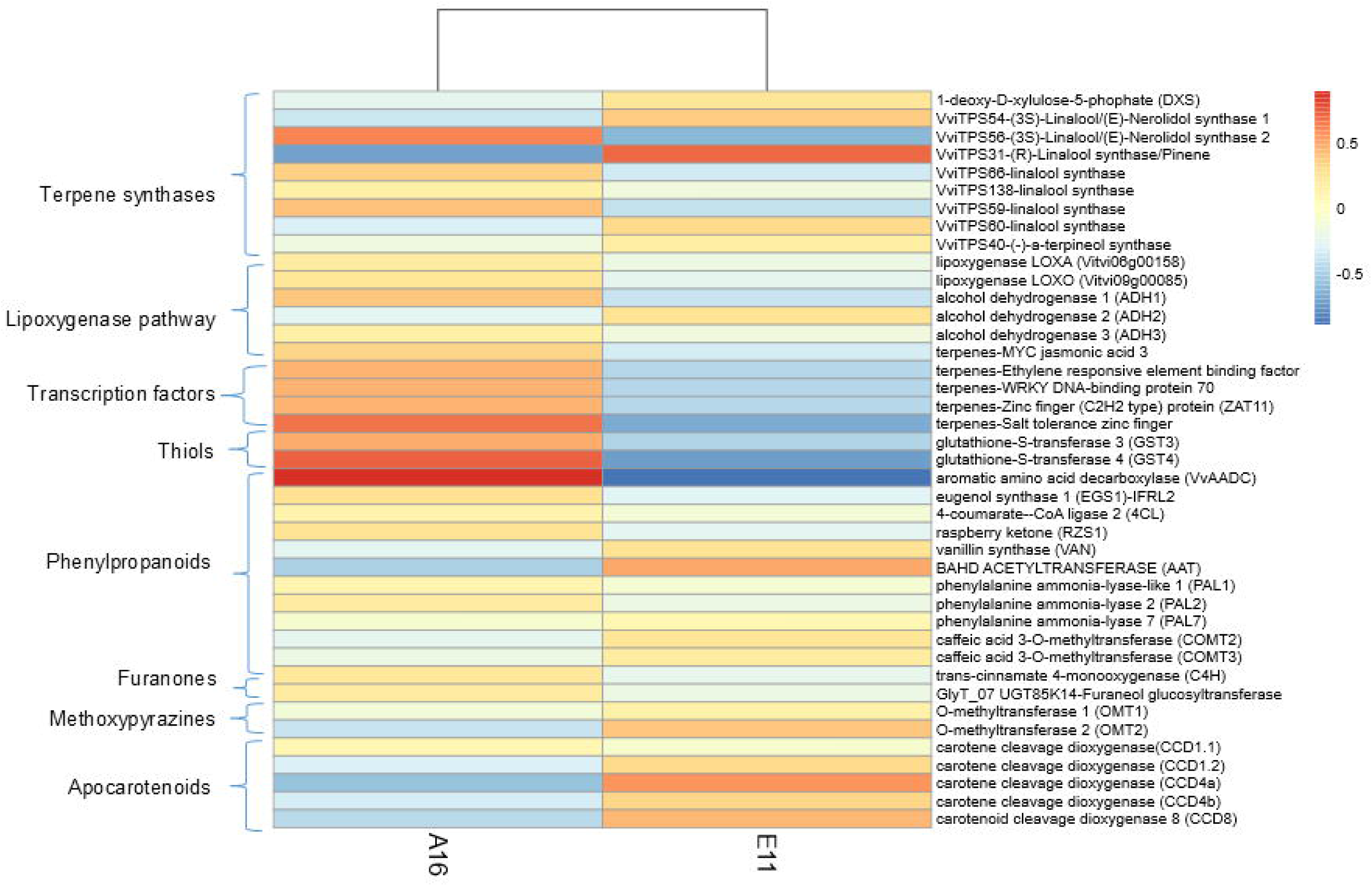
Heatmap of genes related to aroma biosynthetic pathways in Assyrtiko clones A16 and E11, based on the average expression (FPKM) from three biological replicates per clone. Genes are categorized into terpenoid, phenylpropanoid/benzenoid, and volatile phenol biosynthesis pathways. Expression values were log-transformed ln(x + 1)], and Pareto scaling was applied to rows. The heatmap was generated using the ClustVis online tool.

The terpenoid biosynthesis, regulated by two different pathways, was presented in parallel with the expression levels of the genes involved (Fig. 6). The first enzymes of each pathway, 1-deoxy-D-xylulose-5-phophate (DXS) for methylerythritol phosphate pathway (MEP) and acetyl-CoA acetyltransferase (AACT) for mevalonate pathway (MVA), respectively, were both upregulated in clone E11. The same expression exhibited also, the final enzyme of the MVA pathway, farnesyl pyrophosphate synthase (FPPS), which regulates the sesquiterpenes synthesis. On the contrary, the geranylgeranyl pyrophosphate synthase (GGPPS), responsible for the carotenoid production, was upregulated in clone A16.

**Figure 6.**
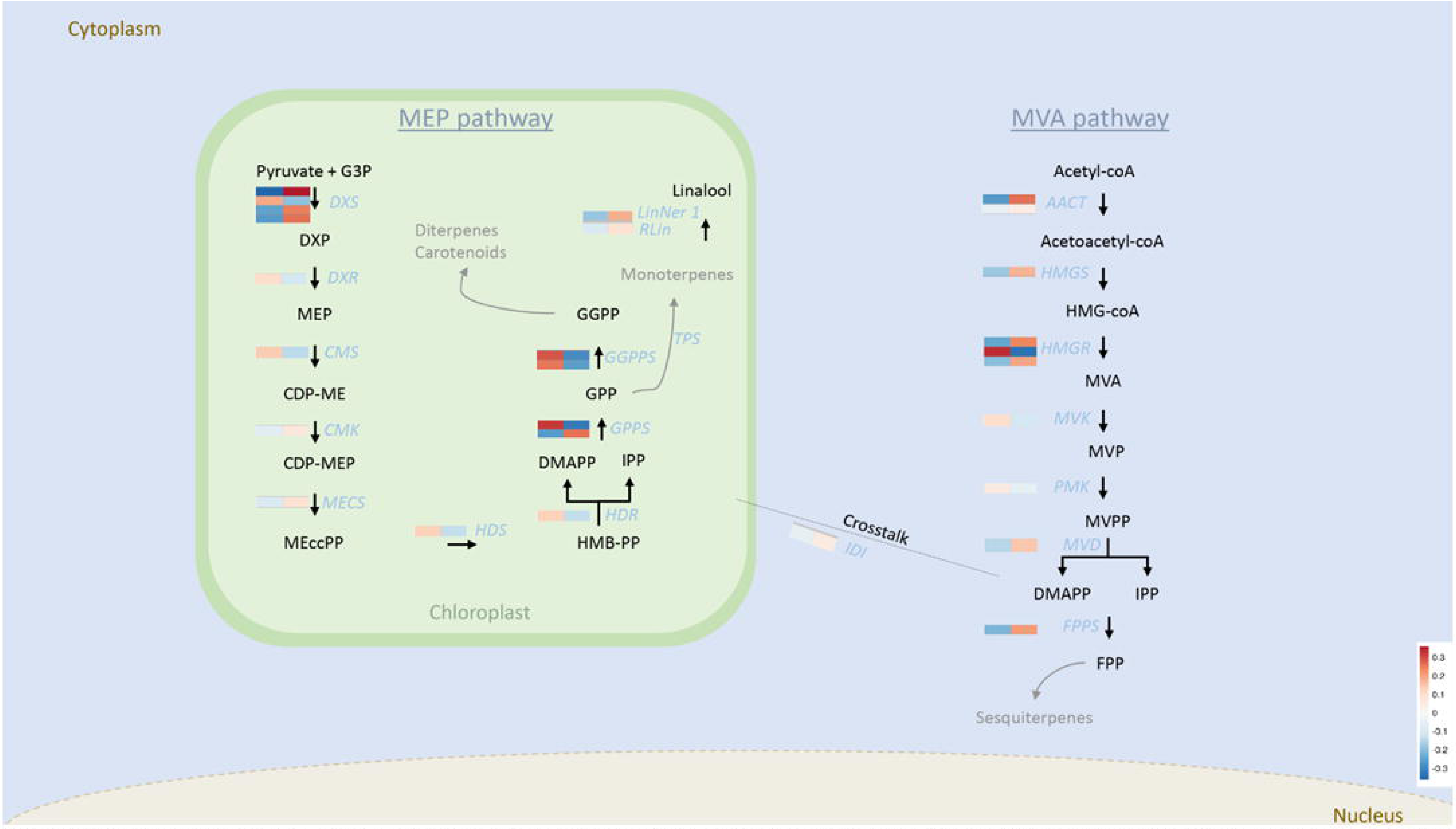
Biosynthetic pathways of terpenoids with heatmap expression of the genes involved, in clones A16 (left) and E11 (right), as FPKM values. *DXS* (Vitvi07g01408, Vitvi04g00438, Vitvi05g00372, Vitvi11g0103): 1-deoxy-D-xylulose-5-phosphate synthase; *DXP*: 1-Deoxy-D-xylulose 5-phosphate; *DXR* (Vitvi17g00816): 1-deoxy-D-xylulose-5-phosphate reductoisomerase; MEP: 2-C-Methyl-D-erythritol 4-phosphate, *CMS* (Vitvi02g00034): 2-C-methyl-D-erythritol 2,4-cyclodiphosphate synthase; *CDP-ME*: 4-(Cytidine 5 ′-diphospho)-2-C-methyl-D-erythritol; *CMK* (Vitvi06g01125): 4-diphosphocytidyl-2-C-methyl-D-erythritol kinase; *MECS* (Vitvi02g00034): 2-C-methyl-D-erythritol 2,4-cyclodiphosphate synthase; *MEcPP*: 2-C-Methyl-D-erythritol 2,4-cyclodiphosphate; *HDS* (Vitvi06g00286): 4-hydroxy-3-methylbut-2-enyl-diphosphate synthase; *HBMPP*: 1-Hydroxy-2-methyl-2-butenyl 4-diphosphate; *HDR* (Vitvi03g00374): 1-Hydroxy-2-methyl-2-(E)-butenyl 4-diphosphate reductase; *GPPS* (Vitvi17g00003, Vitvi06g01196): geranyl diphosphate synthase; *GGPPS* (Vitvi04g01230, Vitvi05g00289): Geranylgeranyl pyrophosphate synthase; *AACT* (Vitvi10g02269, Vitvi12g02504): acetyl-CoA acetyltransferase; *HMGS* (Vitvi02g00420):Hydroxymethylglutaryl-CoA synthase; *HMG-coA*: 3-hydroxy-3-methylglutaryl coenzyme A; *HMGR* (Vitvi18g00043, Vitvi04g01740, Vitvi03g00262): 3-hydroxy-3-methylglutaryl-coenzyme A reductase; MVA: mevalonic acid; *MVK* (Vitvi14g00258):mevalonate kinase; MVP: mevalonate-3-phosphate; *PMK* (Vitvi02g00831): Phosphomevalonate kinase; MVPP: mevalonate-3,5-biphophate; *MVD* (Vitvi13g00887): Mevalonate diphosphate decarboxylase; IPP: Isopentenyl diphosphate; *DMAPP*: Dimethylallyl diphosphate; *IDI* (Vitvi04g01175): isopentenyl-diphosphate Delta-isomerase; *FPPS* (Vitvi19g00729): farnesyl diphosphate synthase; FPP: farnesyl diphosphate.

### Integration of multi-omics data

To identify genes potentially involved in the biosynthesis or regulation of volatile compounds during berry ripening, a Pearson correlation analysis was performed between transcriptomic and metabolomic data from mid-ripe grape berries of two Assyrtiko clones (Supplementary Table S4). Only gene–metabolite pairs with the highest correlation coefficients were retained for network visualization in Cytoscape (Supplementary Fig. S6). Notably, 6-methyl-5-hepten-2-one (HMDB0035915), a volatile with a fruity odor reminiscent of apple and banana, was strongly associated with two genes: ArgJ (arginine biosynthesis bifunctional protein, Gene ID: 100253362), involved in arginine biosynthesis, and IBR3 (probable acyl-CoA dehydrogenase, Gene ID: 100258517), a component of the fatty acid β-oxidation pathway. (Z)-2-Hexenal (HMDB0040264), which emits a green, vegetable-like aroma, correlated with acyl-CoA oxidase 3 (Gene ID: 100257524), a key enzyme in α-linolenic acid metabolism, and β-glucosidase BoGH3B (Gene ID: 100255282), implicated in xyloglucan degradation and short-chain fatty acid production. Hexanal (HMDB0005994), another green-scented compound, was associated with triosephosphate isomerase (Gene ID: 100260576), a central enzyme in glycolysis. 1-Octen-3-one (HMDB0031309), known for its earthy, mushroom-like aroma, exhibited a strong correlation with serine acetyltransferase (Gene ID: 100246391), involved in sulfur amino acid metabolism, and peroxidase 47 (Gene ID: 100261264), a key enzyme in hydrogen peroxide production during plant defense responses. Finally, (E,E)-2,4-heptadienal (HMDB0303844), characterized by a green, oily odor, correlated with glycerol-3-phosphate 2-O-acyltransferase 6 (Gene ID: 100264832), involved in the biosynthesis of glycerophospholipids.

Quantitative real-time PCR (qRT-PCR) was conducted to validate RNA-seq results for selected aroma-related genes (Fig. 7). The analysis included *raspberry ketone synthase 1* (*RZS1*), *eugenol synthase* (*EGS1*), *acetyltransferase* (*AAT*), *vanillin synthase* (*VAN*), *linalool–nerolidol synthase* (*LinNer*), and *wine lactone precursor synthase* (*CYP76F14*). Among these, only *RZS1* showed significantly higher expression in clone A16. *EGS1* and *AAT*, which participates in the same biosynthetic pathway, also exhibited higher expression in A16, although not statistically significant. Overall, the qRT-PCR results were consistent with the RNA-seq data, confirming the reliability of the transcriptomic analysis.

**Figure 7.**
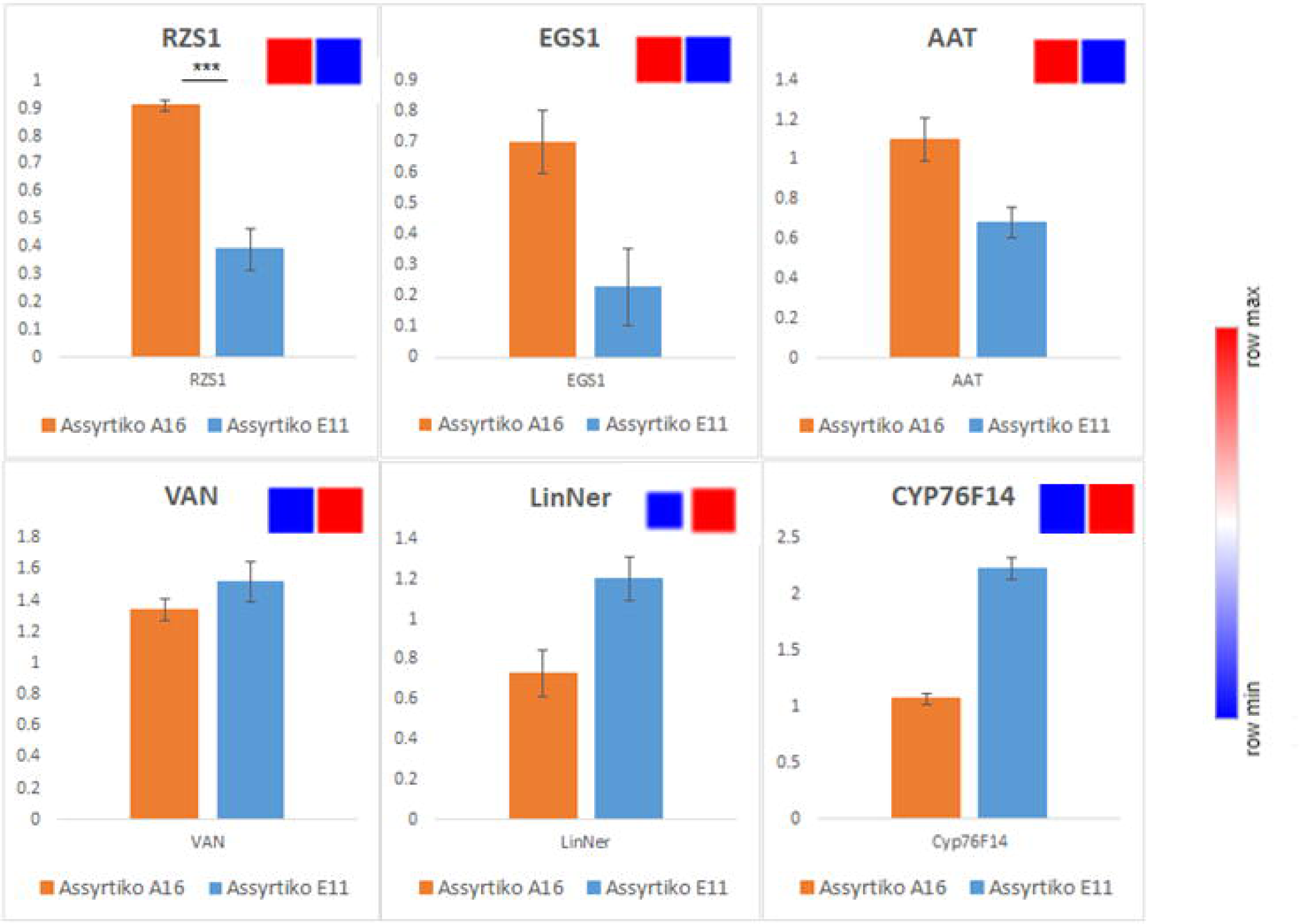
Relative gene expression levels of aroma-related genes in Assyrtiko grapes, as determined by qRT-PCR. The analyzed genes include *RZS1* (raspberry ketone synthase 1, XM_002279390.3), *EGS1* (eugenol synthase, XM_003631652.3), *AAT* (acetyltransferase, XM_002272399), *VAN* (vanillin synthase, XM_002278588.4), *LinNer* (linalool–nerolidol synthase, M8073931.1), and *CYP76F14* (wine lactone precursor synthase, XM_010659734.1). Statistical significance was assessed using a Student’s t-test (*** indicates *p* < 0.01). Heatmap squares in red and blue, generated with Clustergrammer, represent gene expression differences between clones based on RNA-seq FPKM values. Error bars indicate ± standard deviation (n = 3).

### Functional characterization of genes related to the aroma of Assyrtiko Raspberry ketone synthase (RiRZS1)

Raspberry is a characteristic aroma found in various wines. While red wines typically have higher concentrations of raspberry ketone, in a gene expression study of five Greek grape varieties, white grape varieties, such as Assyrtiko and Moschofilero, showed the highest expression (Supplementary Fig. S7). While the gene responsible for raspberry ketone synthesis in *V. vinifera* has not yet been characterized, the biosynthetic pathway is well-established in *Rubus idaeus*. In this pathway, p-coumaroyl-CoA and malonyl-CoA are condensed by the benzalacetone synthase (*BAS*) enzyme to yield 4-hydroxybenzalacetone. This intermediate is subsequently reduced by a NADPH-dependent raspberry ketone synthase (*RiRZS1*) to form 4-(4-hydroxyphenyl)butan-2-one, also known as raspberry ketone (Supplementary Fig 8A). To identify a potential orthologue in *V. vinifera*, the *RiRZS1* sequence was used in a BLAST search, which identified an alkenal reductase (accession no. XM_002279390.3) with 76% sequence similarity (Supplementary Fig 9A). This gene, amplified from the *V. vinifera* Assyrtiko 16 cultivar, was 99.7% identical in nucleotide sequence to the one found in PN40024, but identical in the amino acid sequence, and was cloned into the pUTDH vector using *Bam*HI and *Xba*I restriction sites (Supplementary Fig 9B). The 1,026 bp ORF encodes a 342-amino-acid protein with a predicted molecular weight of ∼37.89 kDa and an isoelectric point (pI) of 5.7.

Predicted 3D structure of the *V. vinifera* alkenal reductase obtained by homology modeling via the RCSB PDB platform. Red spheres represent conserved residues predicted to participate in NADP+ binding or catalysis, based on structural alignment with characterized oxidoreductases. InterProScan further supported the classification of the gene product as an oxidoreductase (Supplementary Fig 8B).

The pUTDH-*VviRZS1* construct was transformed into *S. cerevisiae* strain AM113. Upon supplementation with 0.5 mM benzalacetone as substrate and cultivation for 48–72 hours, ethyl acetate extracts were analyzed by LC-MS. Raspberry ketone was detected in transformed yeast cultures at retention times of 8.3–8.47 min, consistent with the authentic standard, and displayed UV absorbance at 277 nm (Fig. 8B-C). In contrast, no raspberry ketone was detected in cultures transformed with the empty pUTDH vector (Fig 8A), thereby confirming the functional characterization of *VviRZS1* as encoding a raspberry ketone synthase from *V. vinifera* var. Assyrtiko.

**Figure 8.**
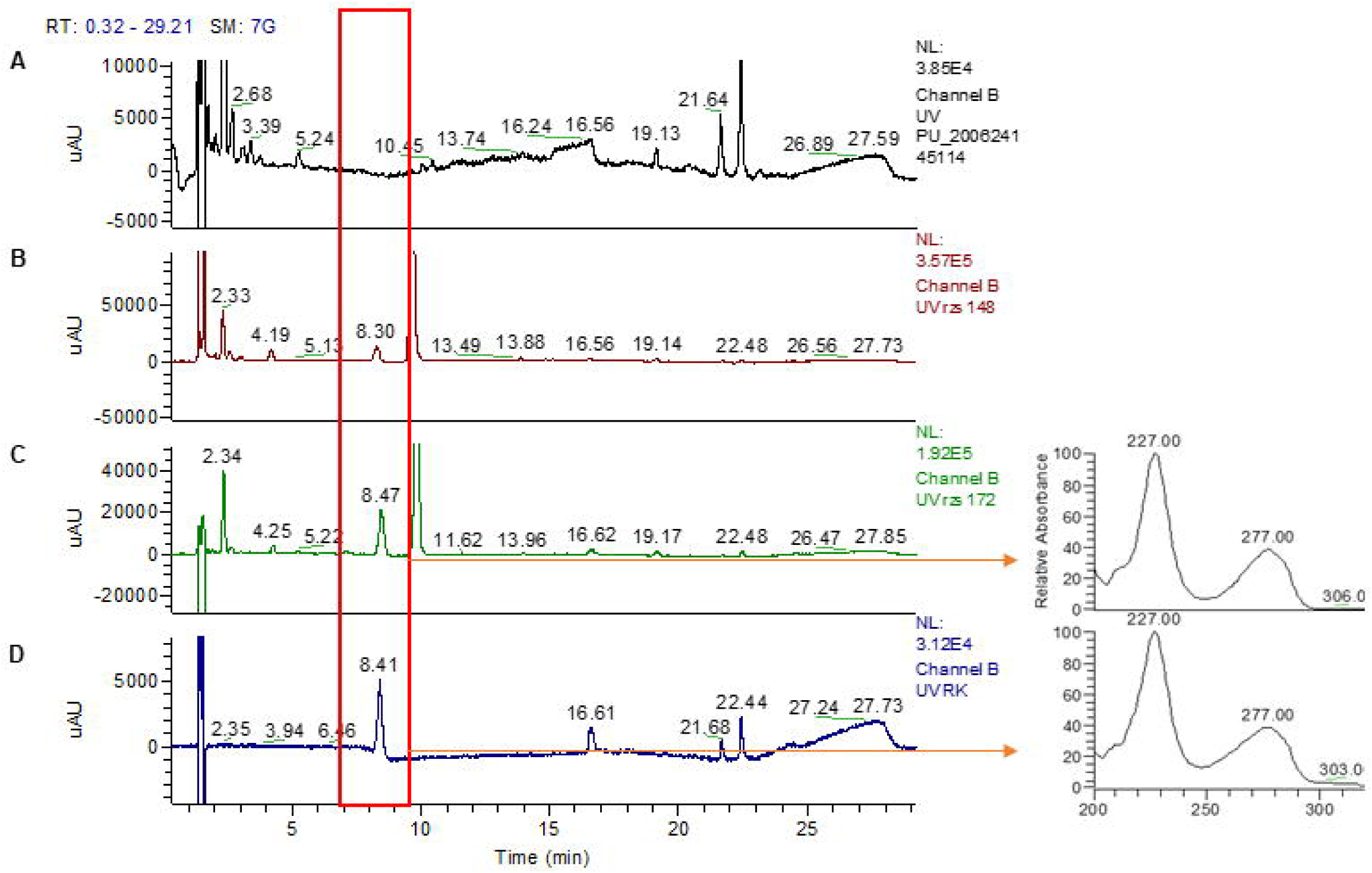
Functional expression of *V. vinifera* raspberry ketone synthase in *Saccharomyces cerevisiae*. LC-MS chromatograms show detection of raspberry ketone in ethyl acetate extracts of transformed *S. cerevisiae* AM113 cultures. A) Control yeast transformed with the empty pUTDH vector, showing no product formation. B) Yeast expressing pUTDH-*VviRZS1* after 48 h of incubation with 0.5 mM benzalacetone. C) Expression after 72 h of incubation. D) Authentic raspberry ketone standard. Raspberry ketone was detected at a retention time of 8.3–8.47 min in samples B and C, matching the standard and confirming *in vivo* activity of the cloned gene. UV absorbance was monitored at 277 nm.

### (E)-8-carboxylinalool synthase

Wine lactone is a volatile compound found in wines, characterized by a sweet, coconut-like aroma. It is formed during wine fermentation through yeast-mediated conversion of its precursor, (E)-8-carboxylinalool. In grape berries, a cytochrome P450 enzyme, *CYP76F14*, catalyzes the sequential hydroxylation and oxidation of linalool (1) to generate (E)-8-hydroxylinalool (2), (E)-8-oxolinalool (3), and (E)-8-carboxylinalool (4), (Ilc et al., 2017). Further, structural modelling of the enzyme revealed a conserved P450 fold, consistent with its catalytic activity (Supplementary Fig. S10A). Sequence comparison of the corresponding gene from two Assyrtiko clones—E11 and 16—with that from Muscat Ottonel and the homologous gene from the *V. vinifera* PN40024 reference genome revealed that clone E11 harbored multiple polymorphisms, whereas clone 16 shared an identical sequence with Muscat Ottonel (Supplementary Fig. S10B). Pairwise identity values among the sequences ranged from 94.58% to 100%, highlighting the evolutionary conservation of *CYP76F14* in diverse grapevine genotypes (Supplementary Fig. S10C). The encoded enzyme showed high sequence homology to the previously characterized *CYP76F14* from Muscat Ottonel and was, thus, selected for heterologous expression in *S. cerevisiae* strain AM113, alongside with a cytochrome P450 reductase (*CPR*). The two genes were co-expressed in yeast supplemented with linalool, and after 72 hours of incubation, cultures were extracted with hexane. Gas chromatography–mass spectrometry (GC–MS) analysis of the hexane extracts identified 8-hydroxylinalool based on library matches and retention characteristics (Fig. 9). This compound is the first expected intermediate in the *CYP76F14*-catalyzed oxidation of linalool, and the only one among the three predicted products that can be detected by GC–MS.

**Figure 9.**
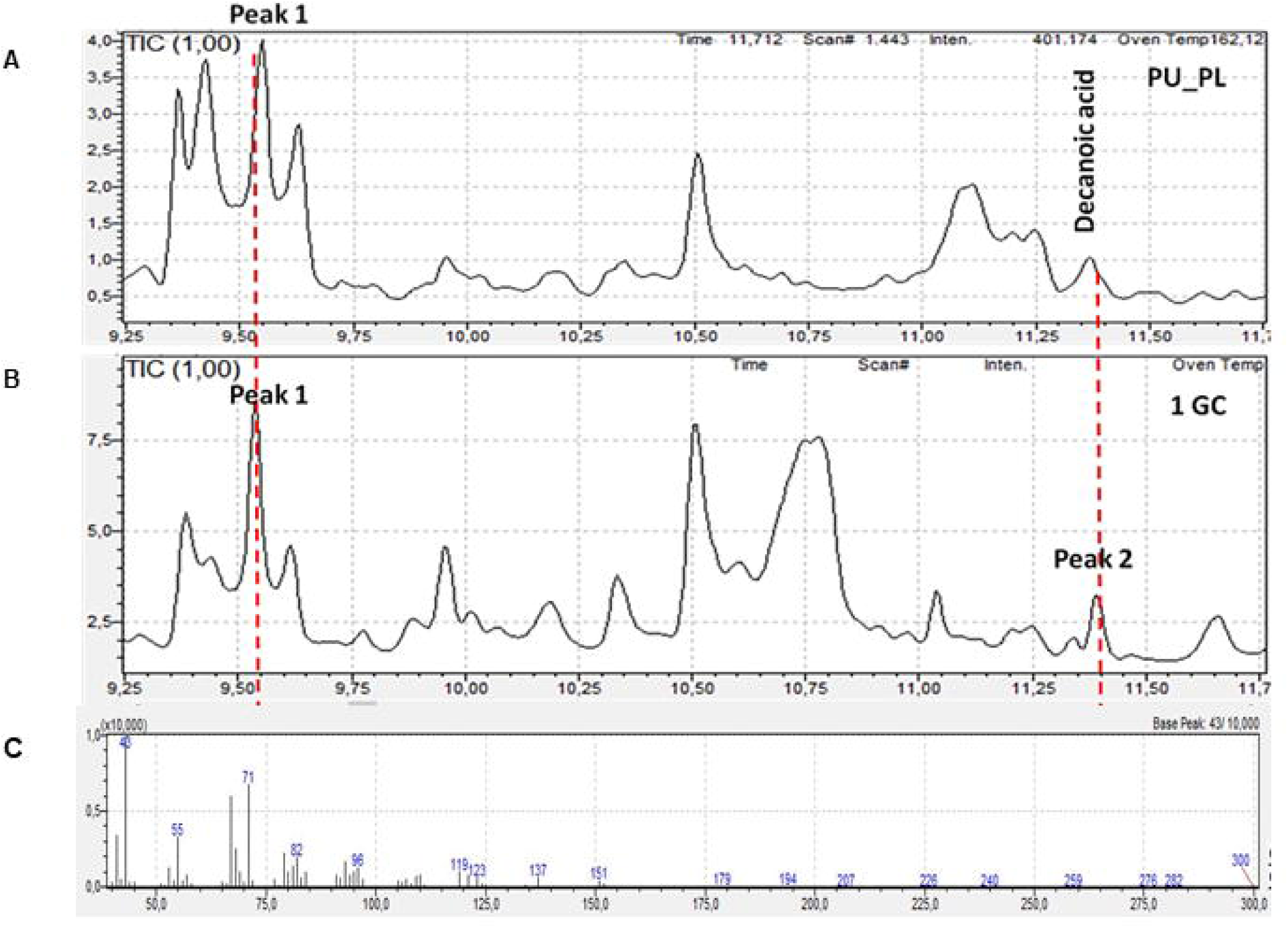
Characterization of (E)-8-hydroxylinalool synthase. GC-MS chromatograms of ethyl acetate extracts from transformed S. cerevisiae strain carrying the pUTDH-*CYP76F14* and pLTDH-*CPR*. A. Control strain harboring empty vectors pUTDH and pLTDH. B. Sample from yeast, transformed with plasmids pUTDH-*CYP76F14* (Assyrtiko 16) and pLTDH-*CPR*. Peak 1: 2,6-dimethyl-7-octen-2,6-diol; Peak 2: 8-hydroxylinalool. C. Main fractions of the peak 2. Three biological replicates with the same construct were used.

### In-vitro characterization of acetyltransferase and eugenol synthase

To further explore the biosynthesis of volatile phenylpropanoids, we investigated the formation of eugenol, a compound derived from coniferyl alcohol. The biosynthetic pathway involves an initial acetylation of coniferyl alcohol (1) to coniferyl acetate (2) by a BAHD-family acetyltransferase (*CAAT*), followed by its reduction to eugenol (3) via eugenol synthase (*EGS1*) (Supplementary Fig. S11A). Homology-based 3D structural models of both EGS1 and *CAAT* were constructed, revealing conserved Rossmann-like folds typical of NADPH-dependent reductases and BAHD acyltransferases, respectively. These models support the predicted enzymatic functions and help explain the observed catalytic efficiency. CLUSTAL alignment of eugenol synthase sequences from *Ocimum basilicum*, Assyrtiko A16, and the PN40024 genome revealed strong conservation, although the A16 variant lacked the N-terminal region, consistent with truncation of the predicted signal peptide (Supplementary Fig. S11B). A similar pattern was observed for the acetyltransferase homologs, with alignments showing high similarity between Assyrtiko A16 and PN40024, and to *O. basilicum* (Supplementary Fig. S11C). Genes of *EGS1* and *AAT* were cloned into pXCK-K vector and expressed in *E. coli* BL21 cells as a fusion protein with an N-terminal DnaK (70kDa) and a C-terminal His tag. It was confirmed by immunoblotting with anti-His antibodies, the expression of the expected fusion protein at ∼106 kDa, corresponding to the 36.05 kDa *EGS1* protein and the 70 kDa DnaK chaperone (Supplementary Fig. S11D) and at ∼120kDa (49,76kDa AAT protein and 70 kDa DnaK chaperone) was soluble (Supplementary Fig. S11E). The purification of both recombinant proteins was performed using HisTrap columns (Cytiva), following solubilization of the insoluble fraction with L-arginine to enhance protein recovery for the insoluble *EGS1* protein. The purified enzymes were tested *in vitro,* incubated with their respective substrate and the resulting reaction mixtures were extracted with ethyl acetate and analysed by GC-MS. When both enzymes were present, a prominent peak corresponding to eugenol was detected (Fig. 10). Compound identification was confirmed by comparison with an authentic standard and spectral matching against mass spectral libraries.

**Figure 10.**
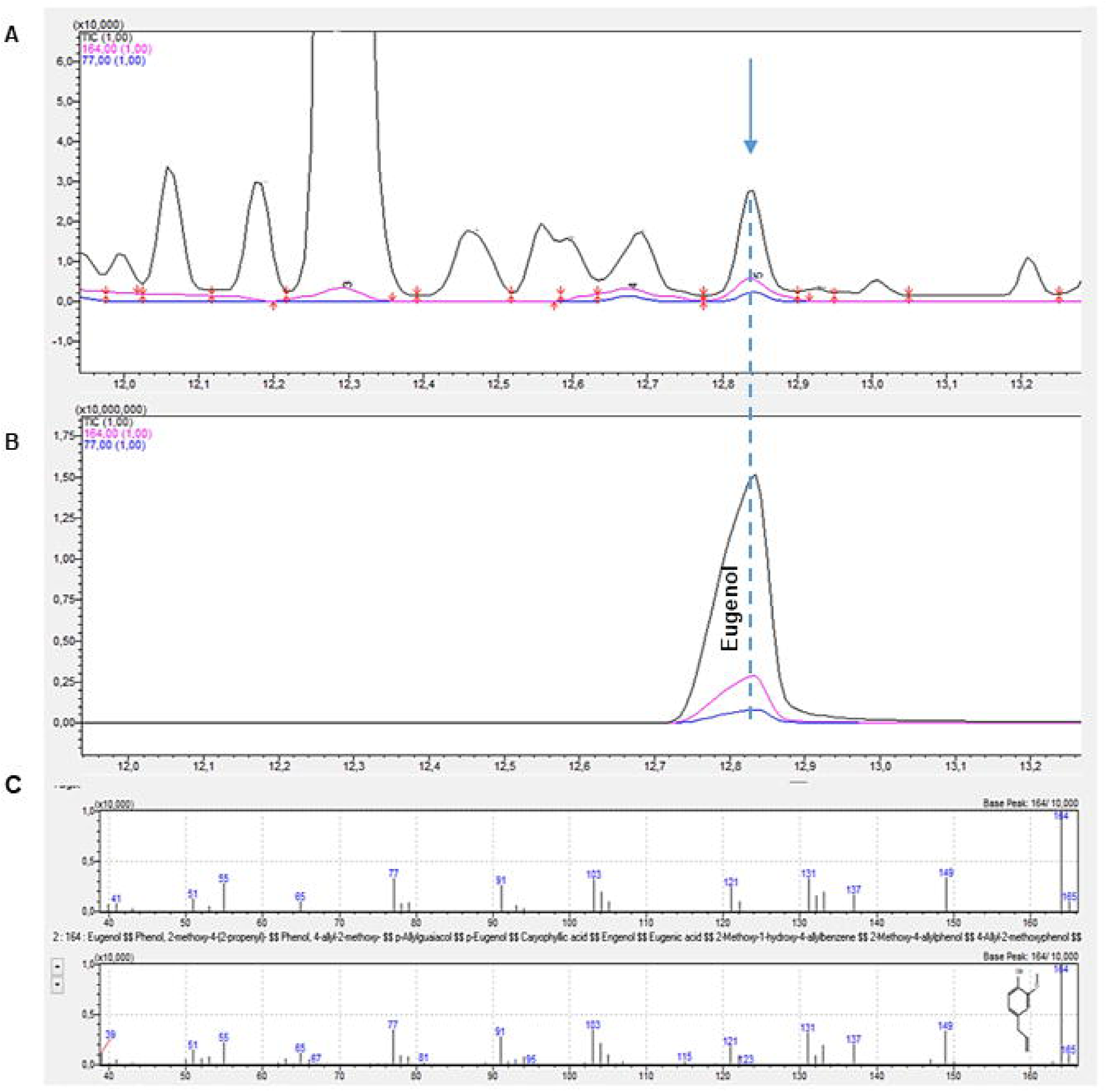
GC-MS analysis of an *in vitro* enzymatic assay confirming eugenol production. A. Reaction with coniferyl alcohol as substrate and both *VvAAT* and *VvEGS1* enzymes. B. Authentic eugenol standard. C. Main fractions of the peak at retention time 12.85 min, identified as eugenol based on NIST mass spectral library matching.

### In vitro characterization of phenylacetaldehyde synthase

A candidate *phenylacetaldehyde synthase* gene was identified in *V. vinifera* through a BLAST search using the functionally characterized gene from *Rosa x hybrida* as a query. Although this gene has been previously reported in the ‘Vidal blanc’ variety, it has not yet been functionally characterized (Pan et al., 2012). The *V. vinifera* PN40024 reference sequence (XM_002266362.4) shares 99.2% nucleotide identity with the ‘Vidal blanc’ sequence, and both Assyrtiko clones analyzed, showed 99.79% identity to PN40024 (Supplementary S12A). A 3D model of the decarboxylase showed a conserved pyridoxal phosphate (PLP)-binding fold characteristic of plant decarboxylases (Supplementary S12B).

The gene was cloned from Assyrtiko 16 into the pAlex expression vector and expressed as a fusion protein with an N-terminal His tag and a C-terminal GST tag. Following purification, the recombinant protein was tested *in vitro* with phenylalanine as the substrate, in the presence of pyridoxal 5′-phosphate (PLP) and NADH. GC–MS analysis confirmed the formation of phenylacetaldehyde in the presence of PLP, indicating that this cofactor is essential for the activity of the grapevine aromatic amino acid decarboxylase (*AADC*) (Supplementary Fig. S13).

## DISCUSSION

The distinctive aromatic profile of wine is a complex trait originating significantly from the specialized metabolism of grape berries (Dixon and Dickinson, 2024), which is influenced by genotype, environment, and viticultural practices (van Leeuwen et al., 2018; González-Barreiro et al., 2015; Ferreira and Lopez, 2019). The distinctive aromatic profile of wine is a complex trait originating significantly from the specialized metabolism of grape berries (Dixon and Dickinson, 2024), which is influenced by genotype, environment, and viticultural practices (van Leeuwen et al., 2018; González-Barreiro et al., 2015; Ferreira and Lopez, 2019). Assyrtiko is well known for its quality *V. vinifera* cultivar, but little is known about the molecular basis behind its distinctive aroma, especially clonal variation (Nanou et al., 2025). To study the biochemical and genetic mechanisms behind the contrasting aromatic profiles between two clones of Assyrtiko, A16 and E11, grown in Nemea, Peloponnese, a multi-omics approach was implemented combining volatile and non-volatile metabolite profiling with transcriptomics (Li et al., 2023), along with functional characterization of key genes to determine the biochemical and genetic basis for their contrasting aromatic profiles. We identified distinct metabolic signatures and corresponding transcriptomic profiles to elucidate the biosynthesis of aroma compounds in this important cultivar during the early ripening stage. This stage of development was selected through relative quantification of aroma-related gene expression (Supplementary Fig. S14). Most of these genes exhibited elevated expression levels at this stage, which led to its selection for subsequent multi-omics analysis.

### Distinct Volatile Profiles Reflect Underlying Transcriptomic Differences

The clear differentiation of clones A16 and E11, demonstrated by their volatile profiles (Fig. 1), underscores their inherent biochemical variations. Clone E11 exhibited notably higher concentrations of the monoterpene linalool (Supplementary Fig. S1), known for its floral and citrus qualities (Mosciano, 1996; Nanou et al., 2025), supporting earlier results and enhancing its distinctive aromatic profile. Additionally, the fragrant characteristics of E11 align with its early-ripening traits, possibly suggesting a quicker transformation of precursors into terpenoid volatiles, thus affecting the aroma profile of the wine (He et al., 2023). Transcriptomic data strongly backs this, showing increased expression of genes related to the methylerythritol phosphate (MEP) pathway, the main pathway for synthesizing monoterpene precursors in plastids (Lichtenthaler, 1999), as well as a linalool– nerolidol synthase (LinNer) identified in clone E11 (Fig. 5). Furthermore, the buildup of various terpenes such as geraniol and nerol, along with C5/C6 alcohols/aldehydes (e.g., trans-2-hexenol) in E11 (Fig. 2B), likely resulting from the lipoxygenase (LOX) pathway through fatty acid precursors (Schwab et al., 2008), aligns with the noted increased expression of an alcohol dehydrogenase 2 (*ADH2*) gene in this clone (Fig. 5). In contrast, clone A16 was noted for its markedly elevated levels of 1-octen-3-one (Supplementary Fig. S1), which added earthy or mushroom-like aromas (Mosciano, 2001; Mendes-Pinto, 2009; González-Barreiro et al., 2015; Nanou et al., 2025), along with the buildup of additional C8 compounds such as 1-octen-3-ol (Fig. 2B). Although the production of C8 volatiles can stem from lipid oxidation, the increased expression of specific lipoxygenase genes (*LOXA*, *LOXO*) in A16 relative to E11 (Fig. 5) implies that clone-specific regulation in the LOX pathway might play a role in these differences.

The elevated concentrations of β-damascenone (a C13-norisoprenoid) in A16 may be associated with the function of carotenoid cleavage dioxygenase (CCD); particularly, many CCDs exhibited greater expression in A16 (Fig. 5), implying a tendency for CCD activities among the clones (e.g., Schwartz et al., 2004; Mendes-Pinto, 2009; Lashbrooke et al., 2013; González-Barreiro et al., 2015; Nanou et al., 2025). These variations in volatile compounds, backed by differential gene expression in relevant pathways (Martin et al., 2012; He et al., 2023), highlight the genetic influences on aroma diversity in the Assyrtiko cultivar.

### Phenylpropanoid Variation and Potential Links to Berry Traits

Although volatile compounds are key factors in determining aroma (He et al., 2023), non-volatile phenylpropanoids greatly influence wine mouthfeel, color, and stability and may act as precursors to aroma compounds (Gutiérrez-Escoba et al., 2021). While no single phenylpropanoid showed statistical significance in univariate analysis, multivariate analysis (PCA, HCA; Fig. 3) and Random Forest modeling (Supplementary Fig. S2C) clearly separated the clones according to their phenolic profiles. Clone E11 displayed higher levels of flavan-3-ols (catechin, epicatechin) and procyanidins, in addition to flavonols such as kaempferol and derivatives of quercetin (Supplementary Fig. S2B), recognized for their antioxidant properties and roles in bitterness and astringency (Huang and Xu, 2021). In contrast, A16 gathered increased levels of hydroxycinnamic acid derivatives (t-coutaric acid) and various flavan-3-ols (gallocatechin, epigallocatechin) (Supplementary Fig. S2B). These patterns generally correspond with the transcriptomic data: KEGG analysis showed enrichment of flavonoid biosynthesis pathway genes among the downregulated genes in A16 (i.e., increased expression in E11), aligning with E11’s greater flavan-3-ol and flavonol levels. The slight yet distinct variations in phenolic makeup may influence not only possible sensory differences in the resulting wines but could also be connected to phenotypic characteristics such as pathogen resistance (Erb and Kliebenstein, 2020), allegedly greater in E11, which also possesses thicker skin, likely associated with flavonoid levels in outer layers (Pourcel et al., 2007)

### Transcriptomic Landscape Reveals Broad Regulatory and Metabolic Reprogramming

The significant differential gene expression found between the two clones (almost 20,000 DEGs under strict criteria, Fig. 4D) suggests considerable differences in their regulatory networks and metabolic functions, reaching well beyond aroma pathways. The increase in A16 of genes linked to specialized metabolite synthesis (covering certain phenylpropanoid, sesquiterpenoid, and diterpenoid pathways), stress responses (such as Cys/Met metabolism), and TF families including NAC, MYB, ERF, and bZIP (Supplementary Fig S3B), acknowledged as key regulators of stress and secondary metabolism (Franco-Zorrilla et al., 2014; Ambrosino et al., 2022), this indicates a distinctive physiological state or adaptive strategy when compared to E11. In contrast, the heightened expression of genes related to plastid/nuclear functions, RNA processing, and transcription in E11, along with the enhancement of flavonoid and monoterpenoid pathways, implies active growth and potentially unique regulation of compounds originating from plastids. The diverse expression of numerous transporters and receptor-like kinases highlights the complex regulatory variations that are crucial (Ward et al., 2009). This broad transcriptomic reprogramming likely affects not only the observed metabolite changes but also numerous other phenotypic distinctions among the clones.

### Functional Validation Confirms Key Enzyme Activities in Assyrtiko

A primary goal was to functionally verify potential candidate enzymes that may enhance Assyrtiko’s aroma profile. We effectively showed that a V. vinifera gene related to Rubus idaeus RZS1 (Koeduka et al., 2011) encodes a functional raspberry ketone synthase (VviRZS1), which converts benzalacetone to raspberry ketone in vivo in yeast (Fig. 8). The significantly increased expression observed in clone A16 is intriguing, despite the fact that raspberry ketone was not identified as a key volatile in our analysis, suggesting that its potential contribution may be undetectable, occur at different developmental stages, or be rapidly transformed. The identified polymorphisms in the E11 allele (Supplementary Fig S9B) require further investigation to understand their possible impacts on enzyme activity.

We additionally validated the role of CYP76F14 from Assyrtiko in producing the wine lactone precursor, identifying the intermediate 8-hydroxylinalool through heterologous expression in yeast supplied with linalool (Fig. 9), aligning with earlier research in Muscat Ottonel (Ilc et al., 2017). Wine lactone imparts coconut-like flavors (Guth, 1997), and comprehending its biosynthesis is essential. Considering that E11 exhibits elevated linalool levels and increased LinNer synthase activity, the role of CYP76F14 (despite its transcript levels not showing significant differences in the DEG list) could result in greater precursor buildup in E11 berries, possibly impacting the wine style. The polymorphisms identified in the E11 CYP76F14 sequence in relation to A16/Muscat Ottonel (Supplementary Fig S10B) might also influence catalytic efficiency.

Furthermore, we validated the activity of enzymes involved in phenylpropanoid volatile synthesis. The biosynthesis of eugenol in plants begins with coniferyl alcohol and involves two key enzymes: an acetyltransferase (QKK36036; AAT) (Dhar et al., 2020), and a eugenol synthase (ABD17321.1; EGS1) (Koeduka et al., 2006) both characterized from sweet basil (Ocimum basilicum). The coupled in vitro activity of Assyrtiko CAAT and EGS1 successfully produced eugenol from coniferyl alcohol (Fig. 10), confirming this pathway operates with grapevine enzymes, analogous to that characterized in Ocimum basilicum (Koeduka et al., 2006). The trend of higher expression of EGS1 transcripts in A16 (Fig. 5) aligns with the KEGG enrichment of phenylpropanoid biosynthesis in this clone, suggesting a higher potential flux through this pathway, even if eugenol was not a major discriminant volatile detected here.

Lastly, we confirmed the PLP-dependent activity of the Assyrtiko aromatic amino acid decarboxylase (AADC) in producing phenylacetaldehyde from phenylalanine (Fig. 11), validating its role in generating potentially floral/honey aroma compounds derived from amino acids (Pichersky et al., 2006), and extending previous identification in ‘Vidal blanc’ (Pan et al., 2012).

### Integrating Omics Data: Towards Understanding Regulatory Networks

The correlation analysis combining transcriptomic and metabolomic data suggested potential connections between certain metabolites and gene expression (Supplementary Table S4). The relationship between the earthy-scented 1-octen-3-one (elevated in A16) and a serine acetyltransferase (which plays a role in sulfur amino acid metabolism) along with a peroxidase (associated with oxidative stress responses) indicates possible connections between lipid oxidation byproducts, sulfur metabolism, and stress responses that require additional exploration (e.g., Chan et al., 2019). In a similar manner, relationships between C_6_ aldehydes (Z)-2-hexenal, hexanal) and genes associated with fatty acid metabolism (acyl-CoA oxidase 3) or central metabolism (TPI) offer testable hypotheses regarding the regulation and production of these green note compounds (Ameye et al., 2018). Though correlated, these combined analyses produce important insights for future functional research focused on revealing the intricate regulatory systems that control volatile production.

### Limitations and Future Directions

Though there are some restrictions to be noted, this study offers a useful synthesis of the molecular variations between Assyrtiko clones A16 and E11. Nemea, one single geographic site, during one growth stage-mid-ripening-served as the test venue. During ripening, the smell profiles of chemicals vary greatly. (El Hadi MAM et al., 2013) and are greatly impacted by terroir (van Leeuwen et al., 2018; Styger et al., 2011). Future research including different developmental stages and diverse surroundings (e.g. comparing Nemea with Santorini) would provide a more dynamic and complete understanding of how genotype-by-environment interactions affect Assyrtiko aroma. Whilst we practically described several significant enzymes, other genes in the related pathways display differential expression (e.g. more CCDs, LOXs, terpene synthases) and warrant more investigation. The correlations in multi-omics are associative and require experimental validation, for instance, through gene editing (e.g. CRISPR/Cas9) or overexpression experiments in grapevine or model organisms (Samaneh Najafi et al., 2023). Examining the functional consequences of the discovered polymorphisms in *RZS1* and *CYP76F14* among the clones would also reveal important insights. Ultimately, investigating the transcriptomic variations associated with stress response (e. g., the link of peroxidase to 1-octen-3-one, varied TF expression) and linking them mechanistically to observed phenotypic differences such as pathogen resistance necessitates further focused research.

## Conclusion

In summary, this study integrated chemical profiling, transcriptomics, and functional enzyme characterization to elucidate the molecular basis for aroma variation between two *Vitis vinifera* cv. Assyrtiko clones and revealed unique volatile and phenolic signatures (enrichment of floral monoterpenes (linalool) in E11 versus earthy C_8_ compounds (1-octen-3-one) and specific C_13_-norisoprenoids in A16), which were associated with differential expression patterns in corresponding biosynthetic pathways (MEP, LOX, potentially CCD) and overall transcriptomic reprogramming of many genes and regulatory factors. We functionally validated the activity of five key aroma-related genes in Assyrtiko: *VviRZS1, CYP76F14, CAAT, EGS1,* and *AADC*. These findings expand our knowledge on aroma biosynthesis in this economically valuable Greek cultivar (Nanou et al., 2025; Alatzas et al., 2021), give candidate genes and metabolites for future breeding programs, and highlight the potential of combining diverse omics approaches for gaining insights into complex traits.

## Author contributions

The study was conceived and designed by AKK. Experiments design and execution of RNA isolation, qPCR, bioinformatic and transcriptomic analyses, selection of genes, cloning, *in vitro* and *in vivo* studies were performed by KL. Assyrtiko clones A16 and E11, plant growth, and grape sampling by KM. Chemical analyses of volatiles and phenylpropanoids were accomplished by UV and CL. GC-MS, LC-MS for functional characterization of genes were executed by ES and AnK. Advising by AK and AKK and funding acquisition by AKK. Data interpretation and manuscript preparation were done by KL and AKK. All authors read and approved of the final manuscript.

## Data Availability

Sequences of the functionally characterized genes from Assyrtiko A16 clone are available from the database of National Center for Biotechnology Information under the accession number ON885732-ON885735, ON885737. Transcriptomic data are deposited into SRA database and can be accessed under BioProject PRJNA973058, BioSamples SAMN35108072-SAMN35108077.

## Abbreviations

OIV: International Organization Of Vine And Wine
VOCs: Volatile Organic Compounds
MVA: Mevalonic Acid Pathway
MEP: Methylerythritol Phosphate Pathway
RNA-Seq: RNA Sequencing
RZS1: Raspberry Ketone Synthase
EGS1: Eugenol Synthase
AAT: BAHD-family Acetyltransferase
AADC: Aromatic Aminoacid Decarboxylase

---

**Supplementary Dataset 1:** Tables S1-S5

**Supplementary Table S1.** Volatile compounds identified by GC-MS analysis of Assyrtiko grape berries, expressed in μg/kg fresh weight.

**Supplementary Table S2.** Compounds identified by LC-MS analysis on Assyrtiko grape berries,

**Supplementary Table S3.** Transcriptomic data from RNA sequencing of Assyrtiko berries at mid-ripening stage.

**Supplementary Table S4**. Pearson correlation analysis between transcriptomic and metabolomic data from mid-ripe grape berries of two Assyrtiko clones.

**Supplementary Table S5.** List of primers designed for cloning specific genes and for relative quantification of gene expression (qRT-PCR).

**Supplementary Dataset 2:** Figures S1-S14

**Supplementary Fig. S1.** Differential accumulation of key volatiles between Assyrtiko clones A16 and E11, by t-test analysis.

**Supplementary Fig. S2.** Correlation and feature selection analyses of phenolic compounds in Assyrtiko grape berries.

**Supplementary Fig. S3.** Transcription factors (TFs) in the differentially expressed genes (DEGs) between Assyrtiko clones A16 and E11.

**Supplementary Fig. S4.** Functional classification of up-and down-regulated differentially expressed genes of A16vsE1, based on Gene Ontology enrichment analysis.

**Supplementary Fig. S5.** Kyoto Encyclopedia of Genes and Genomes (KEGG) pathway enrichment analysis of the up and down-regulated DEGs A16vsE11.

**Supplementary Fig. S6.** Network visualization of transcriptomic and metabolomic correlations in mid-ripe berries of two Assyrtiko clones (A16 and E11).

**Supplementary Fig. S7.** qRT-PCR of RZS1 in two clones from five Greek varieties and FPKM values from RNA sequencing, respectively.

**Supplementary Fig. S8.** Identification and structural characterization of a raspberry ketone synthase candidate from V. vinifera var. Assyrtiko 16.

**Supplementary Fig. S9.** Sequence alignment of raspberry ketone synthase homologues.

**Supplementary Fig. S10.** Identification and structural characterization of wine lactone precursors synthase, catalysed by CYP76F14 from linalool.

**Supplementary Fig. S11.** Identification and structural characterization of a BAHD-family acetyltransferase (CAAT) and eugenol synthase (EGS1), two enzymes involved in eugenol formation.

**Supplementary Fig. S12**. Structural characterization aromatic amino acid decarboxylase (AADC) and CLUSTAL alignment of sequences.

**Supplementary Fig. S13**. GC-MS analysis of an *in vitro* assay verifying phenylacetaldehyde synthesis.

**Supplementary Fig. S14**. Relative quantification of gene expression (qRT-PCR 2018) related to selected aromatic compounds, at four developmental stages of Assyrtiko berries.

